# Dynamic interplay between structural variations and 3D chromosome organization in pancreatic cancer

**DOI:** 10.1101/2021.12.15.471847

**Authors:** Yongxing Du, Zongting Gu, Zongze Li, Zan Yuan, Yue Zhao, Xiaohao Zheng, Xiaochen Bo, Hebing Chen, Chengfeng Wang

**Author notes:** Corresponding author: Chengfeng Wang, State Key Lab of Molecular Oncology & Department of Pancreatic and Gastric Surgery, National Cancer Center/ Cancer Hospital, Chinese Academy of Medical Sciences and Peking Union Medical College, 17 Panjiayuan Nanli, Chaoyang District, Beijing 100021, China. Hebing Chen, Department of Biotechnology, Beijing Institute of Radiation Medicine, Taiping Road 27th, Haidian District, Beijing 100850, China. These authors contributed equally to this work (Yongxing Du, Zongting Gu, Zongze Li).

## Abstract

Structural variations (SVs) are the greatest source of variation in the genome and can lead to oncogenesis. However, the identification and interpretation of SVs in human pancreatic cancer remain largely undefined due to technological limitations. Here, we investigate the spectrum of SVs and three-dimensional (3D) chromatin architecture in human pancreatic ductal epithelial cell carcinogenesis by using state-of-the-art long-read single-molecule real-time (SMRT) and high-throughput chromosome conformation capture (Hi-C) sequencing techniques. We find that the 3D genome organization is remodeled and correlated with gene expressional change. The bulk remodeling effect of cross-boundary SVs in the 3D genome partly depends on intercellular genomic heterogeneity. Meanwhile, contact domains tend to minimize these disrupting effects of SVs within local adjacent genomic regions to maintain overall stability of 3D genome organization. Moreover, our data also demonstrates complex genomic rearrangements involving two key driver genes CDKN2A and SMAD4, and elucidates their influence on cancer-related gene expression from both linear view and 3D perspective. Overall, this study provides a valuable resource and highlights the impact, complexity and dynamicity of the interplay between SVs and 3D genome organization, which further expands our understanding of pathogenesis of SVs in human pancreatic cancer.

## Introduction

Pancreatic cancer is one of the most lethal malignancies worldwide^1^. Over 90% of pancreatic tumors are adenocarcinomas arising from the pancreatic duct epithelium. Therefore, in most cases, “pancreatic cancer” refers to pancreatic ductal adenocarcinoma (PDAC). Because these tumors are highly aggressive and metastatic, unfortunately, most patients are diagnosed at a late stage and do not have the opportunity to receive radical surgery^2^. Even when the primary tumor is small and localized, the prognosis remains poor, and chemotherapy or radiotherapy has limited effectiveness. With a gradually increasing incidence and little improvement in survival rate, PDAC is projected to be the second leading cause of cancer-related death within a decade^3^. Therefore, it seems that a significant improvement in pancreatic cancer mortality relies on the development of earlier detection and better treatment, which requires comprehensive knowledge of the molecular biology and pathogenesis of this disease.

Over the past few decades, the rapid development of next-generation sequencing (NGS) technology (short-read sequencing, SRS) has dramatically expanded our knowledge of the genetic alterations, especially the single-nucleotide variations (SNVs), in PDAC^4^. While some recurrent gene mutations, such as *KRAS*, *CDKN2A*, *TP53* and *SMAD4,* have been demonstrated to successively initiate and drive PDAC progression^5^, the spectrum and pathogenesis of larger structural variations (SVs) in the context of pancreatic ductal epithelial cell carcinogenesis remain largely undefined due to technological limitations. SVs, including insertions, deletions, duplications, inversions and translocations at least 50 bp in size, are the structural and quantitative chromosomal rearrangements that constitute the majority of genetic differences across human genomes^6^. Accumulating evidence has demonstrated that SVs contribute to polymorphic variation, pathogenetic conditions and many human diseases, such as cancers^7, 8^. For a long time, SV detection has been performed through SRS approaches, which have been reported to lack sensitivity, exhibit a very high false positive rate and misinterpret complex or nested SVs^9, 10^. Recently, long-read methods, referred to as third-generation sequencing (TGS) technologies, have been developed and shown to produce genome assemblies of unprecedented quality^11^. The first true representative of TGS is single-molecule real-time (SMRT) sequencing, developed by Pacific Biosciences (PacBio) ^12^. With average read lengths of 10 kbp or higher, reads can be more confidently aligned to the repetitive sequences that often mediate the formation of SVs. In addition, long reads are more likely to span SV breakpoints with high-confidence alignments. Due to these advantages, long-read single-molecule sequencing has the potential to substantially increase the reliability and resolution of SV detection. Using long-read SMRT technology, Nattestad *et al*. sequenced a breast cancer cell line and developed one of the most detailed maps of SVs in a cancer genome available; this map included nearly 20,000 variants, most of which were missed by short-read sequencing^13^. Similarly, Aganezov *et al.* found hundreds of variants within known cancer-related genes that were detectable only through long-read sequencing^14^. These findings highlight the need for third-generation/long-read sequencing of cancer genomes for the precise analysis of structural variant signatures to understand the molecular etiology underlying these diseases^15, 16^.

High-throughput sequencing technologies can dramatically accelerate the discovery and characterization of SVs; however, the medical interpretation of SVs and the prediction of phenotypic consequences remain crucial challenges for geneticists and cancer biologists^17^. Obviously, SVs that disrupt or include coding sequences can directly alter gene expression through their effects on gene dosage^18^. Nonetheless, the discovery that SVs can be pathogenic without changing coding sequences indicated that SVs can have regulatory effects by influencing the position and/or function of cis-regulatory elements, such as promoters and enhancers, that is, position effects^19^. Owing to advances in 3D genome mapping technologies, such as high-throughput chromosome conformation capture (Hi-C) sequencing, it is now becoming increasingly evident that position effects are the result of alterations that are much more complex than simple changes in the linear genome and can be understood only by considering the three-dimensional organization of the genome^20^. Recent findings have indicated that dynamic changes in 3D genome architecture are associated with the development of multiple malignancies^3–5, 9–11, 13–15, 21–30^, including breast cancer, multiple myeloma, B cell lymphoma, and T cell acute lymphoblastic leukemia, by coordinating the expression of some key driver genes. Notably, SVs can disrupt 3D genome organization and thereby exert direct regulatory effects on gene expression^31^. Meanwhile, the occurrence and formation of genomic rearrangements can be influenced by the 3D chromatin architecture^32^, highlighting an action-reaction interplay between SVs and the 3D genome.

In light of these reports, understanding how SVs contribute to cancer pathogenesis by interplaying with chromatin organization remains largely unexplored. Herein, to broadly assess the global structural variation spectrum and 3D chromatin architecture in pancreatic cancer, we performed SMRT and in situ Hi-C sequencing in two primary pancreatic ductal carcinoma cell lines (PANC1 and BxPC3) and an immortalized normal epithelial cell line (HPDE6C7). We further integrated the resulting datasets with transcriptomics by RNA-seq to elucidate the influences of dynamic interplay between SVs and 3D genome organization on gene expression. Moreover, some public datasets including enhancer activity, expressional level and clinical prognosis were also acquired to validate above regulatory correlation or biological significance (Fig. 1a). Our study provides fundamental new insights into the genetic and molecular basis of PDAC development and may contribute to the discovery of novel potential targets or biomarkers for precision therapy.

**Fig. 1.**
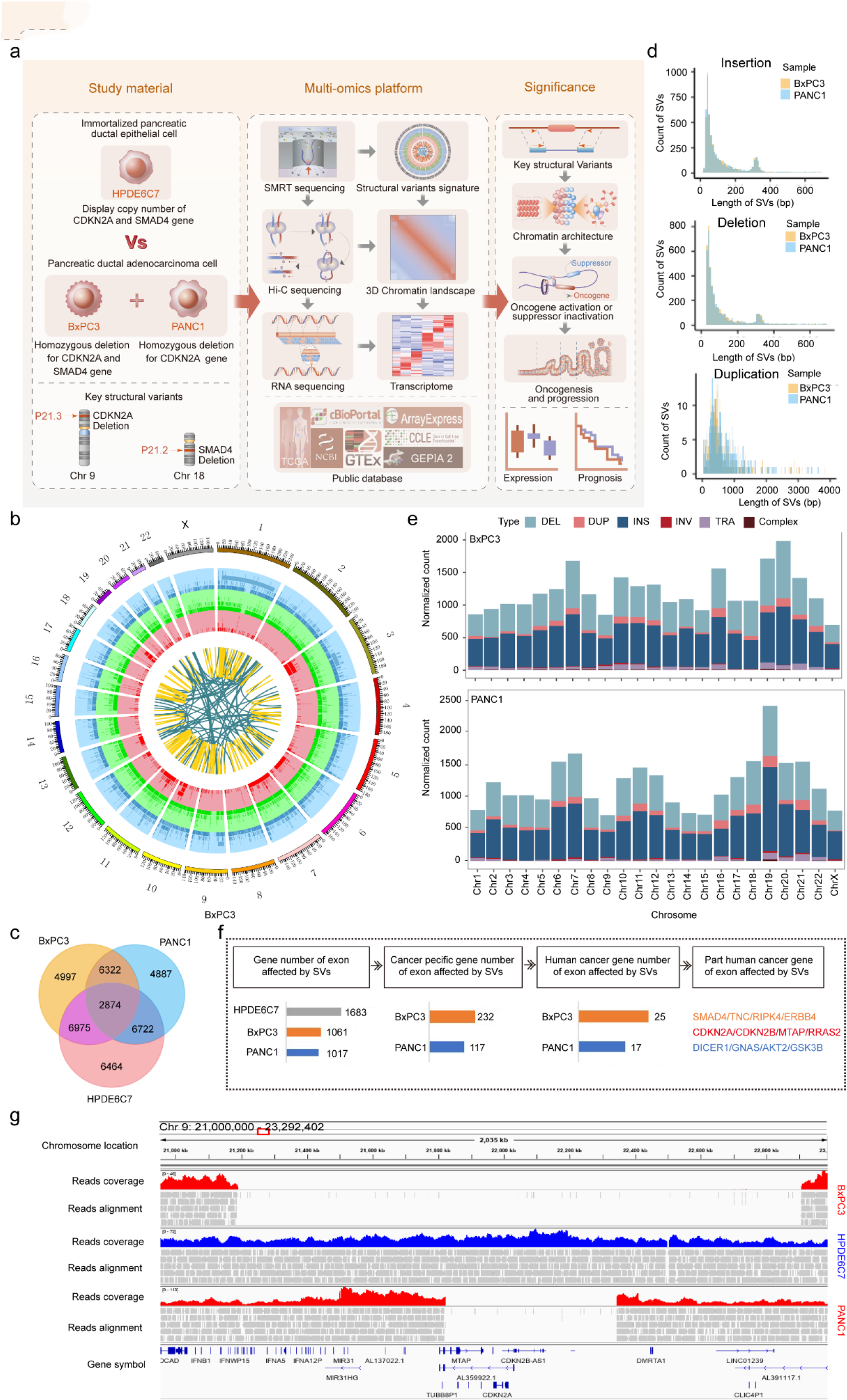
The overall landscape of SVs in PANC1, BxPC3 and HPDE6C7. **a**, Schematic diagram of the overall research design with the study material, multiomics platform, and significance presented. **b**, Circos plot showing the high-confidence SVs detected by Sniffles in BxPC3 with 23 chromosomes inputted. The tracks from the outer to the inner circles are the chromosome coordinates, deletions, insertions, duplications, inversions and translocations. **c**, Venn diagram showing the intersection of structural variations in two cancer cell lines (PANC1 and BxPC3) and one normal pancreatic ductal epithelial cell line (HPDE6C7) with counts indicated. **d**, Histograms show the length distribution of specific SVs found by Sniffles, suggesting similar size distributions for insertions, deletions and duplications in PANC1 and BXPC3. **e**, Distribution of standardized total SV burden (deletion: light blue, duplication: pink, insertion: deep blue, inversion: red, translocation: light purple, complex: brown) across chromosomes. **f**, Pipeline of identification of specific cancer-related genes directly affected by SVs in exonic regions in PANC1 and BxPC3. **g**, IGV image showing a homozygous deletion on chromosome 9 (covering CDKN2A, CDKN2B and MTAP) of different lengths in BxPC3 and PANC1.

## Results

### The signatures of structural variations in human pancreatic cancer

We first comprehensively investigated the dynamic spectrums of SV s that occur during malignant transformation of normal pancreatic ductal epithelial cells by SMRT sequencing. The structural variation data, processed by our SMRT-bench platform, showed good alignment rates with a high percentage of usable long-range read pairs (Supplementary Fig. 1 and Supplementary Table 1). By mapping to the reference genome xCRCh37/hg19, a large number of SVs were detected, with total counts of 20805, 21168, and 23035 SVs in PANC1, BxPC3 and HPDE6C7, respectively (Fig. 1b, Supplementary Fig. 2a and Supplementary Table 2). The two most common types of SVs were insertions and deletions, which accounted for approximately 50% and 41% of all SVs, respectively. Notably, the number of complex SVs in which more than two different types of simple SVs break-ends overlapped was elevated two- to fourfold in cancer cell lines compared with normal epithelial cell line, which indicated that genome instability increased greatly during malignant transformation. Next, we explored the distribution of SVs in different regions of the genome and found that a majority of them were located in intergenic and intronic regions, as reported in previous studies^16, 20, 21^ (Supplementary Fig. 2b).

To further identify the specific SVs that might lead to tumorigenesis, we compared all the SVs detected in cancer cells with those detected in normal epithelial cells and found that more than half of the SVs were specific to BxPC3 or PANC1 (Fig. 2c and Supplementary Table 3). The number of common SVs shared by the three cell lines was 2874, which accounted for only 13% of all SVs detected. These findings suggested that the SV signature had strong cell type specificity. Additionally, we studied the length distributions of different specific SV types. Insertions, deletions and duplications showed similar characteristics, and most were within 1 kb in length (Fig. 1d), while inversions and complex SVs were relatively different between the two cancer cell lines (Supplementary Fig. 2c). In addition, we analyzed the standardized counts and proportions of different specific SV types within each chromosome. Although the normalized number of total SVs in each chromosome was slightly different, the proportion of each specific SV type was basically similar (Fig. 2e).

**Fig. 2.**
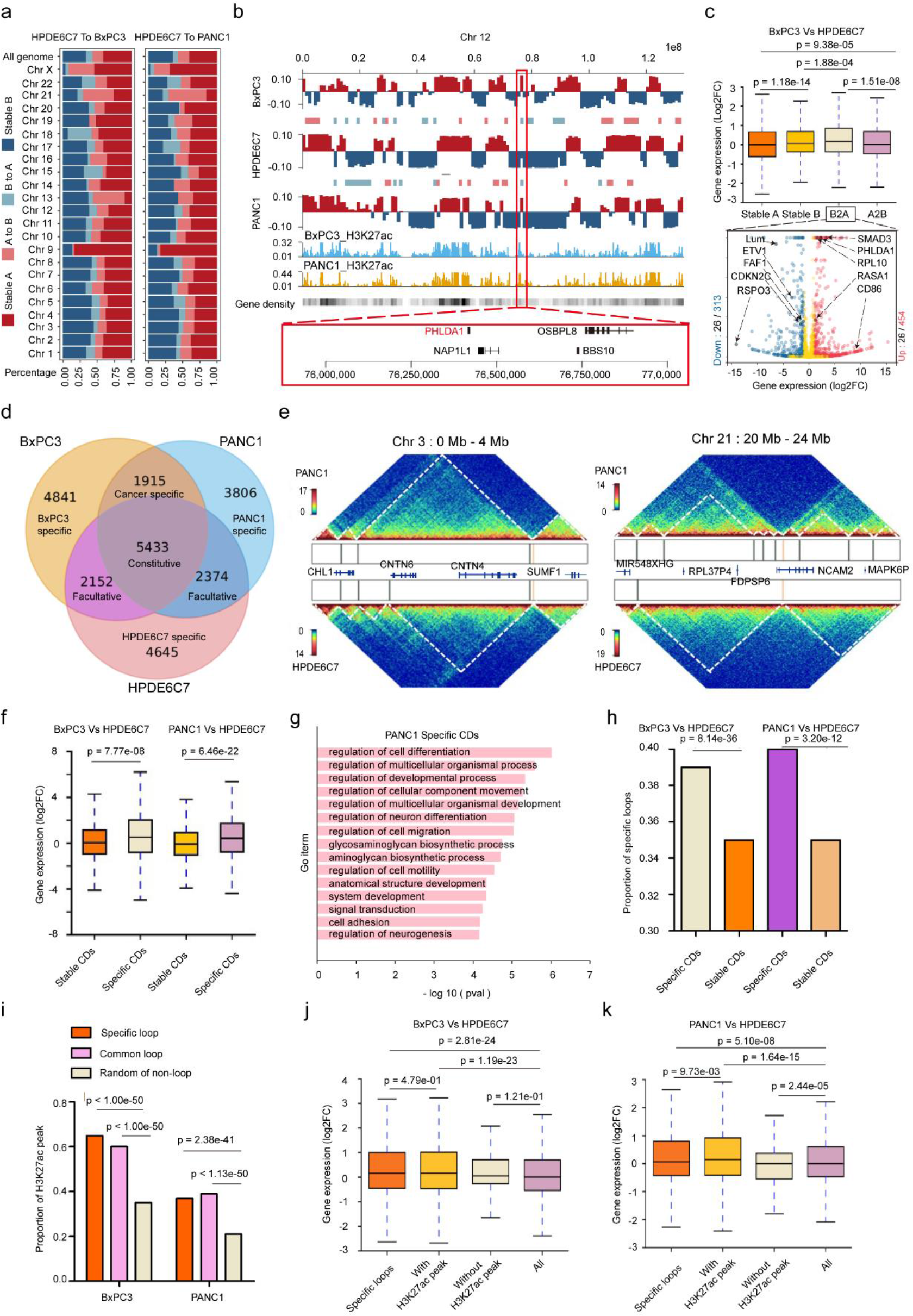
3D chromatin architecture remodeling correlates with gene expression changes in human PDAC. **a**, Compartment A/B and compartment switching of whole chromosomes in PANC1 and BxPC3 compared with HPDE6C7. Assignment of the A compartment (deep red) and B compartment (deep blue) were performed using eigenvalues > 0 and < 0, respectively. **b**, Examples of A/B compartment shifts on chromosome 12 in PANC1 and BxPC3 compared with HPDE6C7. Roadmap epigenome enhancer activity, marked by H3K27ac signal peaks, in PANC1 and BxPC3 is shown as blue and brown in histograms. Columns show the gene density in the genome. A red frame denotes a common B-to-A in both cancer cell lines covering the PHLDA1 gene. **c**, Top: Box plots showing the comparison of gene expression levels in different compartments between BxPC3 and HPDE6C7. The box represents the interquartile range (IQR), the centerline denotes the median, and the whiskers extend to 1.5 times the IQR (or to the maximum/minimum if < 1.5 × IQR). Bottom: Volcano plots showing the number of differentially expressed genes (blue) and cancer-related genes (black) among them in the B-to-A shift region. Genes indicated by the black arrow are examples of significantly upregulated (red on the right) or downregulated (blue on the left) cancer-related genes (|Log2FC|>1 and adjusted p value<0.05). Gene expression was compared as Log2FC (BxPC3/HPDE6C7) with the P-value obtained by the Wilcoxon rank-sum test. **d**, Venn diagram showing that the majority of domain boundaries that are present in normal cells (HPDE6C7) are also present in cancer cells (constitutive boundaries; PANC1 and BxPC3), while 15∼20% of boundaries were maintained in only one of the two cancer cell types (cell-type-specific boundaries). The number of CDBs was obtained from the interaction matrices at 10-kb resolution in the three cell types. **e**, Examples of CD alterations in regions of interest (left chr 3: 0-4 Mb, right chr 21: 20-24 Mb) in PANC1 compared with HPDE6C7. The vertical bars in the box between heatmaps represent CDBs. Genes involved in the region are indicated. Compared with HPDE6C7, PANC1 shows different patterns of changes in the number and length of CDs in different chromosome regions. **f**, Box plots represent gene expression levels in different types of CDs in BXPC3 and PANC1 compared with HPDE6C7. Specific CDs were defined as CDs with specific boundaries identified by HiCDB. Gene expression was compared as Log2FC (cancer cell line/HPDE6C7) with the P-value obtained by the Wilcoxon rank-sum test. **g**, Biological process enrichment of differentially expressed genes located in specific CDs of PANC1. P values were obtained from Fisher’s exact test using EnrichR. GO, gene ontology. **h**, Histograms represent proportions of specific loops in different types of CDs in BxPC3 and PANC1 compared with HPDE6C7. Cell-specific loops were defined as loops for which the observed/expected ratio of two ends of the loop was twice that of the control sample. P-values were calculated by chi-square test. **i**, Histograms represent the proportions of H3K27ac peaks in different types of loops in the genomes of BxPC3 and PANC1. P- values were calculated by chi-square tests. **j**, **k**, Box plots represent the gene expression level in specific loops with or without H3K27ac peaks comparing BXPC3 and PANC1 with HPDE6C7. Gene expression was compared as Log2FC (cancer cell line/HPDE6C7) with the P-value obtained by the Wilcoxon rank-sum test.

Next, we further investigated the genes that were directly affected by SVs in their exon regions and obtained 1017, 1061 and 1683 genes in PANC1, BxPC3 and HPDE6C7, respectively (Fig. 1f, and Supplementary Table 4). The specific genes affected by SVs in cancer cell lines were 177 and 232, of which 17 and 25 were reported as human cancer genes (HCGs) in the Catalogue of Somatic Mutations in Cancer (COSMIC, see URLs) ^33, 34^ (Supplementary Table 5 and 6). Notably, the specific HCGs shared by both cancer cell lines included *CDKN2A*, *CDKN2B* and *MTAP* deletions on chromosome 9 and *RRAS2* insertions on chromosome 11. In addition, *DICER1* amplification on chromosome 14, *AKT2* deletion on chromosome 19 and *GNAS* insertion on chromosome 20 were independently detected in PANC1. *TNC* deletion on chromosome 9, *RRAS2* insertion on chromosome 11 and *SMAD4* deletion on chromosome 18 were specifically detected in BxPC3. *CDKN2A* homozygous deletion was demonstrated in PANC1 and BxPC3 but not HPDE6C7 in Cellosaurus, a knowledge resource for cell lines (see URLs, Supplementary Table 7). To confirm these findings in our data, the relevant genomes in three cell lines were visualized by IGV (Integrative Genomics Viewer, see URLs). There was an obvious large deletion in BxPC3 with a length of more than 1.7 Mb involving many genes in addition to *CDKN2A*, including *CDKN2B*, *MTAP* and *DMRTA1* (Fig. 1g). Similarly, a smaller deletion of approximately 0.5 Mb that simultaneously covered the *CDKN2A*, *CDKN2B* and *MTAP* genes was found in PANC1. Interestingly, we found another duplication in the region adjacent to this deletion in PANC1, which could also be verified by next-generation sequencing data from the Yue Feng lab^21^ (Supplementary Fig. 2e). In addition, a homozygous deletion of *SMAD4* was observed in BxPC3, which was consistent with data from Cellosaurus (Supplementary Fig. 2d). These findings further supported the identity of our cell lines and the reliability of SV data generated using the SMRT-bench platform in this study. The above SVs directly affected oncogenes or tumor suppressors in cancer cells and thus might play a significant role in the oncogenesis and maintenance of the malignant phenotype.

Together, these state-of-the-art long-read sequencing results establish signatures of SVs in human pancreatic cancer, which should provide a valuable resource for comprehensive investigation of the pathogenesis of SVs in this deadly malignancy.

### The 3D chromatin architecture is remodeled and correlates with gene expression changes in human PDAC

Previous studies have shown that SVs may reshape chromatin folding and alter the location of cis-regulatory elements such as promoters or enhancers leading to tumorigenesis^20, 35^. To further reveal the impact of SVs on 3D chromatin architecture and gene expression, we applied in situ Hi-C sequencing to comprehensively analyze the spatial conformation of chromosomes in the normal HPDE6C7 cell line and two cancer cell lines (BxPC3 and PANC1). Correlation analysis of the primary reads of three cell lines from different libraries indicated that Hi-C data from different libraries were consistent and that the two cancer cell types were most similar to each other and could be distinguished from the normal HPDE6C7 cell line (Supplementary Fig. 3a and Supplementary Table 8). Next, we compared the 3D genome architecture of cancer cell lines with that of a normal epithelial cell line on multiple scales.

### Cis and trans interactions in PDAC

We first analyzed the genome-wide interaction ratio of each chromosome in the three cell lines and found that most of the interactions were intrachromosomal, whereas only a few interactions were interchromosomal (Supplementary Fig. 3b and 3C).. Next, we analyzed the correlation between chromosome size, spatial position and interchromosomal interactions. We found that small, gene-rich chromosomes (chromosomes 16, 17, 19, 20, 21, 22) preferentially interacted with each other, suggesting that chromosome proximity strongly influenced contact probability (Supplementary Fig. 3d), which is consistent with previous studies^22, 36, 37^. In addition, Circos^38^ analysis revealed a large number of novel or lost trans-interactions in both cancer cell lines (BxPC3 and PANC1) compared to the normal HPDE6C7 cell line (Supplementary Fig. 3e), suggesting that changes in trans-interactions, despite occurring at relatively low proportions, might play a role in tumorigenesis.

**Fig. 3.**
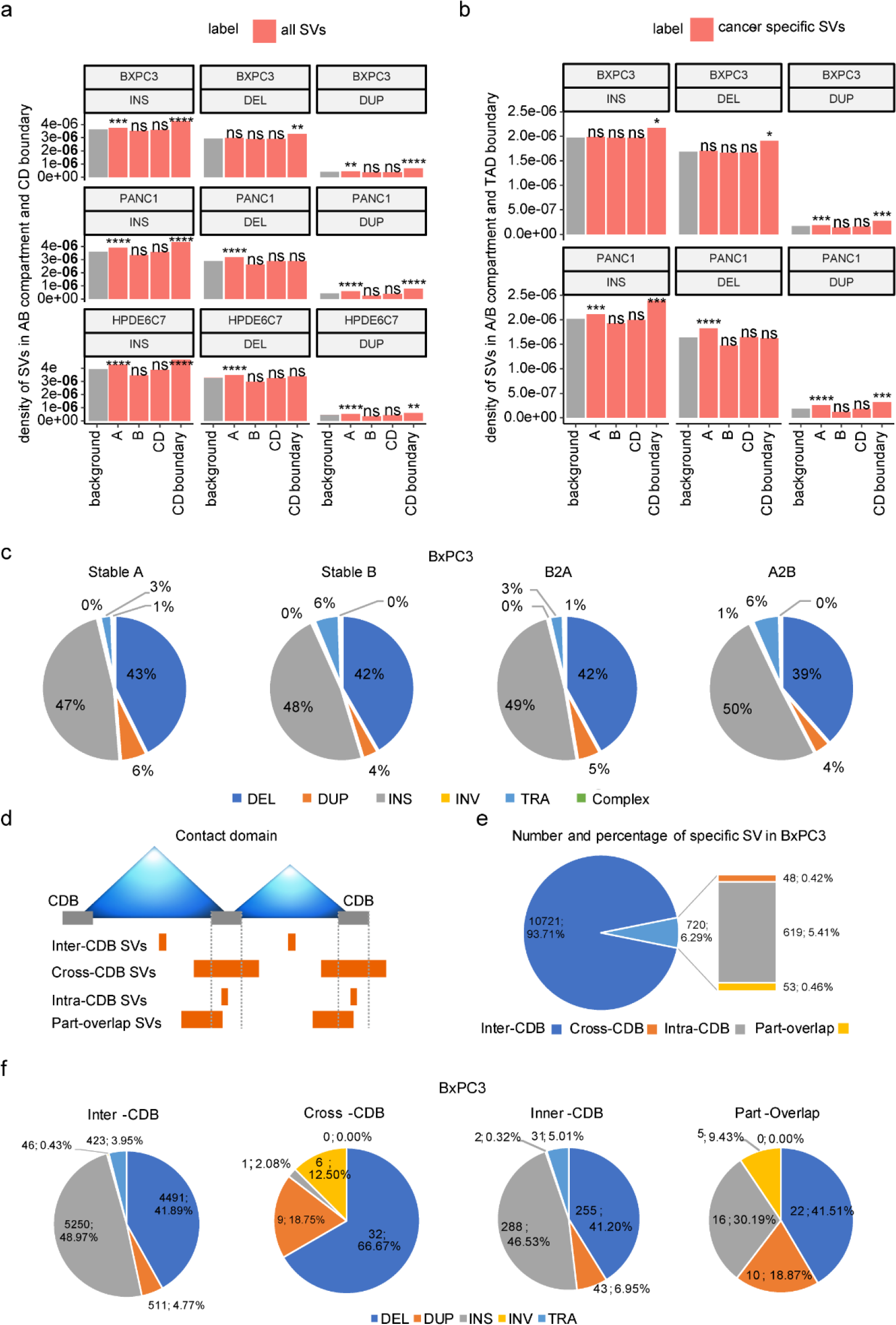
Distributions of structural variations among 3D genome architectures. **a**, Density of SVs (insertions, deletions and duplications) in A/B compartments and contact domain boundaries (CDBs). The density of SVs is the number of SVs divided by the length of each chromosome region. Gray bars represent background sequence and show the density of SVs on the chromosomes. Red bars show the density of SVs on the different chromosome regions, including A/B compartments, CDs, and CDBs. Enrichment tests were performed via R’s prop. test, which was evaluated by comparing the proportion of SVs falling in the region of interest and the proportion of the length of the region of interest in the whole genome. ****p ≤ 0.0001, ***p ≤ 0.001, **p ≤ 0.01, *p ≤ 0.05. **b**, The density of cancer-specific SVs (insertions, deletions and duplications) in A/B compartments and TAD boundary. Cancer-specific SVs refer to those occurring in BxPC3 (or PANC1) but not in HPDE6C7. Enrichment tests were evaluated by comparing the proportion of SVs falling in the region of interest and the proportion of the length of the region of interest in the whole genome. ****p ≤ 0.0001, ***p ≤ 0.001, **p ≤ 0.01, *p ≤ 0.05. **c**, The proportion of different types of cancer-specific SVs (less than 2 Mb length) in different A/B compartment regions of BxPC3. **d**, Schematic diagram of SV categories according to the position relationship between the breakpoint of cancer-specific SVs and the CD boundary. **e**, Number and percentage of four different specific SV types in BxPC3 according to **d**. **f**, The proportion of different types of four categories of cancer-specific SVs in BxPC3 according to **d**.

### Compartment switching in PDAC

In general, human chromosomes can be divided into two compartments, A and B. Compartment A usually consists of open and gene-rich chromatin regions, whereas compartment B consists of gene-poor and closed chromatin regions; these regions are spatially separated, and can be converted into each other^39^. Genome compartmentalization signatures have been identified in previous cancer studies^23^, with compartment A/B transition presenting in up to 20% of genomic regions in breast cancer, multiple myeloma, B-cell lymphoma, and T-cell acute lymphoblastic leukemia^24, 26–28^. We next performed an analysis of compartment A/B transition in the genome and found that compared with HPDE6C7, the overall incidence of A/B switching was 24.8% in BxPC3 (A-to-B, 15.3%; B-to-A, 9.5%) and 24.1% in PANC1 (A-to-B, 16.2%; B-to-A, 7.9%), respectively. Although the percentage of A/B switching appeared different on each chromosome, the majority of the genome exhibited a stable A/B identity. Stable A was clearly the most common while A/B transition was the least part on chromosome 9 in both cancer cell lines (Fig. 2a and Supplementary Table 8).

Additionally, A/B switching were reported to be associated with changes in gene density and regulatory activity. Our data showed that stable A and B-to-A compartments were gene-rich regions with active transcription, while stable B and A-to-B compartments had the opposite properties (taking chromosome 12 as an example in Fig. 2b). Further genome-wide comparative analysis identified common and specific compartment shifts in each PDAC cell line compared with HPDE6C7. The proportions of stable A and B in the genomes of BxPC3 and PANC1 were 33.03% and 29.39%, respectively, and the common A-to-B and B-to-A ratios were 7.8% and 3.53%, respectively (Supplementary Fig. 4a). Interestingly, the specific A-to-B and B-to-A switching in BxPC3 cells accounted for 7.45% and 6% of all genome, respectively. Correspondingly, specific A-to-B and B-to-A transitions occurred in 8.46% and 4.33% of the genome in PANC1, respectively, suggesting the cell type specificity of A/B switching. Further analysis in combination with RNA-seq data revealed that compartment A/B and compartment switching were significantly associated with gene expression changes; i.e., the gene expression in stable A and B-to-A compartments was significantly higher than that in stable B and A-to-B compartments (Fig. 2c and Supplementary Fig. 4b), which is in line with previous reports^26, 27^. Next, we analyzed the features of differentially expressed genes (|Log2FC| >1 and q-value <0.05) and found that the number of differentially expressed genes in the A-to-B compartment was significantly higher than that in the B-to-A compartment in BxPC3/PANC1 cells (1291/1314 vs. 767/696, P<0.05) (Fig. 2c, Supplementary Fig. 4b/c/d and Supplementary Table 10). We further aligned with the reported human cancer related genes and found that *PHLDA1*, a gene located in the common B-to-A compartment of chromosome 12, was significantly upregulated in both cancer cell lines (BxPC3:Log2FC 2.33, q-value < 0.001; PANC1: Log2FC 1.94, q-value < 0.001). Data from the GEPIA2 (Gene Expression Profiling Interactive Analysis) server show that *PHLDA1* encodes a proline–histidine-rich nuclear protein that may play a critical role in the antiapoptotic effects of insulin-like growth factor; moreover, its expression is significantly upregulated in several malignancies, including pancreatic cancer, lower-grade glioma, and melanoma^40^, and it is significantly associated with poor prognosis in pancreatic cancer.

Together, these data show that the spatial distribution of chromatin compartments A and B are changed in two cancer cell lines (BxPC3 and PANC1) compared with HPDE6C7, and these transitions were significantly associated with expression changes in cancer-related genes.

### Contact domain alterations in PDAC

The topologically associating domain (TAD) is the functional unit of chromatin architecture. In previous studies, TADs were mainly identified by insulation score ^24, 26, 41^, but this method could identify only large topological units due to technical drawbacks leading to low resolution ^42^. To better characterize the contact domains (CDs) of normal and cancer cell lines, we employed high-resolution Hi-C, and contact domain boundaries (CDBs) were detected using the HiCDB method based on local relative insulation metrics and a multiscale aggregation approach on Hi-C maps^43^. We identified 14581, 14318 and 13494 CDs in the HPDE6C7, BxPC3 and PANC1 cell lines with average sizes of 211 kb, 227 kb and 214 kb (Supplementary Table 11), respectively. Notably, most constitutive CDBs were shared by HPDE6C7, BxPC3 and PANC1 cells (21.59%), followed by cell-specific CDBs (18.46%, 19.24% and 15.12% for HPDE6C7, BxPC3 and PANC1, respectively). In contrast, few facultative CDBs were shared between any two cell types (8.55%, 7.61% and 9.43% for HPDE6C7 vs. BxPC3, BxPC3 vs. PANC1 and PANC1 vs. HPDE6C7, respectively) (Fig. 2d and Supplementary Table 12). These results suggested that CDBs are conserved in the human genome.

For some cancer types (breast and prostate cancers, multiple myeloma), it has been reported that acquisition of new CD boundaries is usually accompanied by a corresponding increase in CD number and decrease in CD size^24, 26, 27^. However, our study showed that disappearance of CD boundaries was more typical in pancreatic cancer cells (Supplementary Fig. 4e and f). Notably, changes in CD number and size could present diametrically opposite alterations in different chromosome regions. For example, on the Hi-C map, fewer and longer CDs were observed in chromosome 3 of PANC1 than in the same region of HPDE6C7, while a region of chromosome 21 showed the opposite trend (Fig. 2e); the same finding was also observed in BxPC3 (Supplementary Fig. 4g). Similar results have been observed in gliomas and acute lymphoblastic leukemias^25, 44^, suggesting that the trend of CD alterations in cancer is not absolute and that CDs may present quite diverse changes in different cancer types. Restrictive assumptions, parameter selection, normalization or matrix correction made by different computational methods may be the main reasons for the differences in these results^45^. Cancer heterogeneity and features specific to immortalized HPDE6C7 cells may also be contributing factors. To further explore the relationship between changes in different types of CDs and altered gene expression in the corresponding regions, we defined the newly emerging CDs in cancer cells as cancer-specific CDs and the CDs shared with HPDE6C7 as stable CDs. We found that cancer-specific CD regions were significantly more associated with upregulated gene expression compared with stable CDs (Fig. 2f and Supplementary Table 13). Further Gene Ontology (GO) analysis of differentially expressed genes in cancer-specific CDs revealed that altered genes were involved in several key pathways, including cancer promotion, cell differentiation, cell adhesion, cell motility and migration (Fig. 2g and Supplementary Fig. 4g).

### Cancer-specific loops and aberrant enhancer activations in PDAC

To explore the relationship between differential gene expression and loops in chromatin, we profiled a total of 4046 and 1859 cancer-specific loops in BxPC3 and PANC1, respectively. Chromatin loops were identified using Hi-C Computational Unbiased Peak Search (HiCCUPS). If the observed/expected ratio of the two ends of the loops in the cancer cell line was twice as high as that in the normal control cell line, the loops were defined as cancer-specific loops. The numbers of cancer-specific loops with H3K27ac peaks were 3743 in BxPC3 and 1357 in PANC1 (Supplementary Table 14).

Further analysis of CDs revealed that cancer-specific CDs were significantly associated with a high proportion of cancer-specific loops (Fig. 2h). Moreover, the proportion of active enhancers marked by H3K27ac was higher in cancer-specific loops than in common loops and random nonloop regions in the genome (Fig. 2i) and was significantly associated with upregulated gene expression (Fig. 2j and k). These data suggested that the upregulation of gene expression in cancer-specific CDs might be related to the activation of enhancers in cancer-specific loops. Therefore, compared with normal HPDE6C7, spatial chromatin CDs are altered in cancer cell lines. Moreover, these changes are accompanied by upregulation of gene expression and may be related to the enhanced activity of regulatory elements in cancer-specific loops.

In conclusion, the 3D chromatin architecture in PDAC cell lines has undergone widespread remodeling and consequent dysregulation of gene expression, which may promote tumorigenesis and progression of PDAC.

### Distributions of structural variations among 3D genome architectures

Previous studies have indicated that the occurrence and formation of SVs are affected by 3D chromosome organization ^32^. Therefore, we next studied the distribution of SVs at the level of the 3D genome organization and analyzed the three most common types of SVs: insertions, deletions and duplications.

We first explore the relationship between SVs and A/B compartments. Insertions, deletions and duplications were all significantly enriched in the A compartment of BxPC3/PANC1/HPDE6C7 cells, except for deletions in BxPC3 (Fig. 3a). It has been demonstrated that the A compartment is transcriptionally active^22^ and that SVs tend to occur in the A compartment. Meanwhile, it was reported that SVs usually occurred at the 3D loop-anchors which commonly led to compartment transition^32^. We compared the A/B compartments of HPDE6C7 with those of BxPC3 or PANC1 and found that insertions, deletions and duplications were significantly enriched among the stable A of BxPC3 and PANC1 (Supplementary Fig. 5a). In addition, insertions, deletions and duplications were significantly enriched in the A-to-B compartment in BxPC3, while deletions and duplications were significantly enriched in the B-to-A compartment in PANC1 (Supplementary Fig. 5a), which indicated that the occurrence of SVs may be related to A/B compartment conversion. The differences in the distribution pattern among cell lines strongly suggest the cell type specificity of the correlation between SVs and A/B compartment transition.

Furthermore, we selected cancer-specific SVs from all SVs to study their distribution pattern in 3D chromatin architecture. Interestingly, we found that insertions, deletions and duplications were significantly enriched in the A compartment in PANC1, but only duplications were enriched in the A compartment in BxPC3 (Fig. 3b). In terms of the dynamic compartment transitions, insertions were significantly enriched in the A-to-B compartment in BxPC3, while they were only significantly enriched in the stable A compartment in PANC1; deletions were enriched in the stable A and A-to-B compartments in BxPC3, while they were enriched only in the stable A compartment in PANC1; the distribution of duplications was similar in BxPC3 and PANC1; and all duplications were enriched in the stable A compartment (Supplementary Fig. 5b). The above findings again indicated that the distribution patterns of SVs in A/B compartments were different between the two cell lines. Cancer-specific insertion and deletion tended to occur in dynamically changing A-to-B compartments in BxPC3, while both tended to occur in stable A compartments in PANC1. We further analyzed the compositions of different types of cancer-specific SVs less than 2 Mb length in A/B compartments in BxPC3 and PANC1. Insertions, deletions and duplications accounted for approximately similar proportions in stable A, stable B, A-to-B, and B-to-A compartments in the two cancer cell lines (Fig. 3c, Supplementary Fig. 5c, and Supplementary Table 14 and 15).

Next, we investigated the distributions of SVs in CDs and their boundaries. For all SVs, insertions and duplications were significantly enriched in the CD boundaries in BxPC3/PANC1/HPDE6C7, while deletions were enriched in the CD boundaries only in BxPC3 (Fig. 3a and Supplementary Fig. 5d). These findings indicated that most SVs tended to occur near the boundaries of CDs, consistent with previous studies^26, 46^. We further classified CD boundaries by comparing HPDE6C7 with BxPC3 and PANC1 and found that insertions and duplications were significantly enriched only in the gained CD boundary in PANC1 but not in BxPC3 (Supplementary Fig. 5e). Similarly, we observed the same phenomenon in cancer-specific SVs (Supplementary Fig. 5f and g), which indicated the significant cell type specificity of the degree of SV enrichment in CD boundaries.

Then, all cancer-specific SVs were divided into four categories: Inter-CDB/Cross- CDB/Inner-CDB/Part-overlap according to the relative position between the breakpoint of SVs and CDB (Fig. 3d). We found that more than 90% of cancer-specific SVs were located inside the CDs (inter-CDB-SVs), which had little impact on their organization, while Cross-CDB/Inner-CDB/Part-overlap SVs, which had a greater probability of influence on CD folding, accounted for a relatively low proportion (Fig. 3e and Supplementary Fig. 5h). We further analyzed the compositions of different types of cancer-specific SVs in the Inter-CDB/Cross-CDB/Inner-CDB/Part-overlap groups. The proportions of different types of cancer-specific SVs in BxPC3 and PANC1 were roughly similar in the Inter-CDB/Inner-CDB/Part-overlap groups, while for the Cross- CDB group, the percentages of different types of SVs differed greatly (Fig. 3f, Supplementary Fig. 5i and Supplementary Table 17).

In conclusion, there is a certain correlation between the occurrence of SVs and 3D chromosome organization in tumors at the A/B compartment or CD level. Furthermore, to some extent, the distribution pattern of SVs is different at the 3D genome level in different tumor cells, which indicates that the distribution of SVs among the 3D genome is highly cell-type specific.

### Cancer-specific SVs affect gene regulation via reshaping CDs in PDAC genomes

Previous studies have revealed that SVs can rewire chromatin organization to alter chromatin topologies and gene regulation in *cis*^20, 35, 47^. CDs are specific functional units that are blocked by CDBs, mediated by CTCF and other insulators. Theoretically, cross- CDB SVs may disrupt chromatin folding domains and create new regulatory units, thereby altering gene expression far from SV breakpoints and triggering disease^42, 48, 49^. To explore the impact of cross-CDB SVs on CD disruption, we analyzed the correlation between cross-CDB deletion and CD fusion and found that CD fusion was significantly more frequent in the cross-CDB deletion regions than in other sites of the genome (Fig. 4a and Supplementary Table 18). These results indicate that cross-CDB deletion is significantly associated with CD fusion, which is consistent with previous research results^20, 29, 50^. However, not all cross-CDB deletions could cause CD fusion, and further analysis found that only deletions with higher frequency in the same cell line were associated with enhanced interaction of adjacent CDs or CD fusion. Conversely, deletions with lower frequency had no significant effect on the interactions of adjacent CDs, and no CD fusion was identified in these cases (Fig. 4b/c and Supplementary Fig. 6a/b). Here, frequency refers to the percentage of SVs in the whole detected cell population. Obviously, higher frequency of deletions means less interaction in the relevant SV region on Hi-C heatmaps. Similarly, cross-CDB duplications did not always result in increased CD interactions; a significant increase in CD interactions or formation of neo-CDs was observed at only a small number of cross-CDB duplications that were homozygous or of higher copy number (Supplementary Fig. 7a and b). These results indicate that the effects of SVs on the 3D genome architecture are quite complicated and may be influenced by multiple factors, such as the intercellular genomic heterogeneity, location, and length of the SVs^46, 51^.

**Fig. 4.**
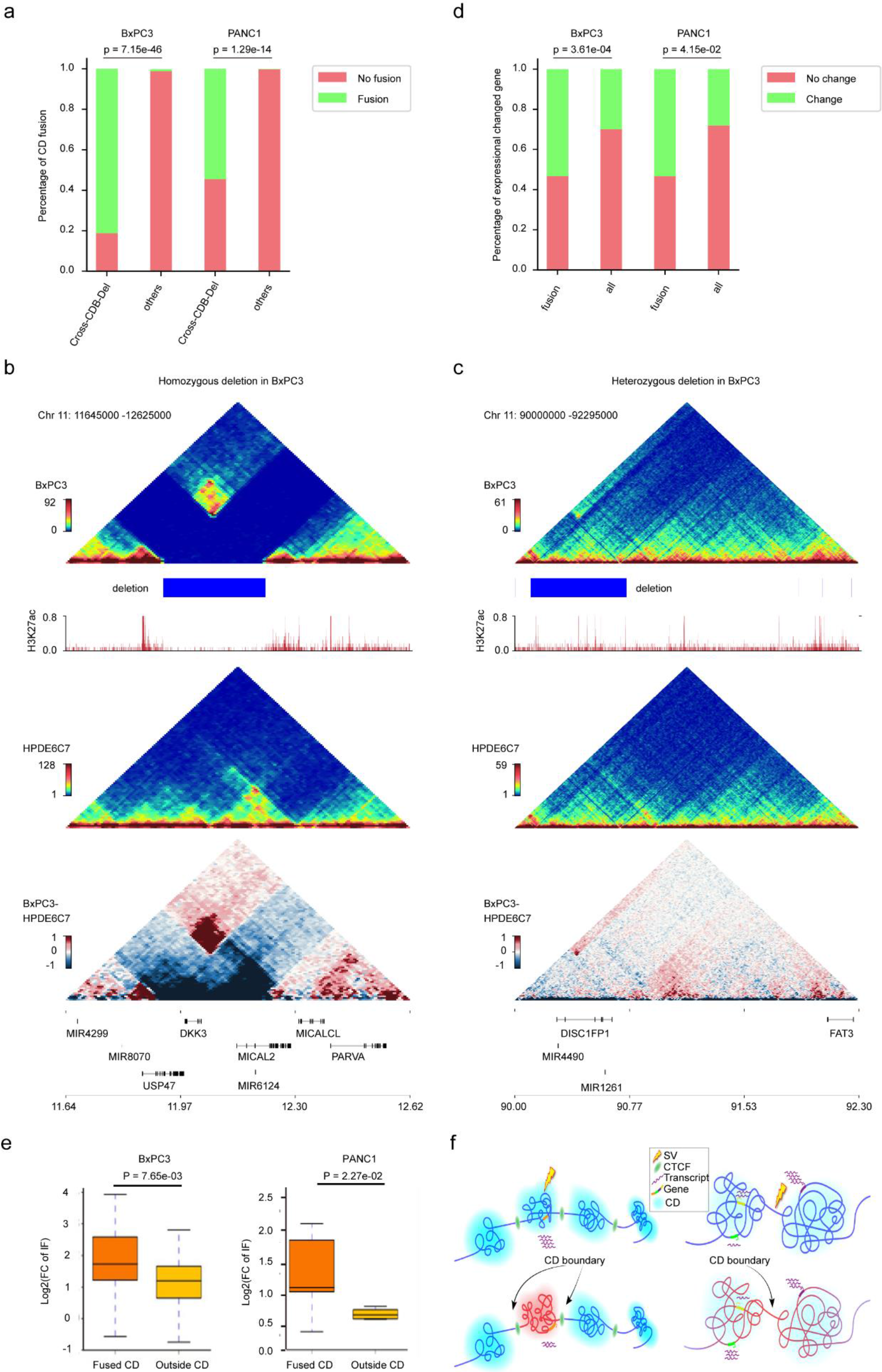
Cancer-specific SVs affect gene regulation via reshaping CDs in PDAC genomes. **a**, Proportion of CD fusions in cross-CDB deletion fields and other genome regions in BXPC3 and PANC1. P-values were calculated by fisher’s exact test. **b**, **c**, Examples of the impact of cross-CDB deletion purity on the chromatin folding domain in BXPC3. Triangle heatmaps represent chromatin contact frequency, with the top showing BxPC3, middle showing HPDE6C7, and bottom showing the subtractive results. Histogram representing roadmap epigenome enhancer activity, marked by H3K27ac, in BxPC3 (red). **b**, Homozygous cross-CDB deletion is associated with CD fusion. **c**, No significant enhancement of adjacent CD interactions was observed at heterozygous cross-CDB deletion. **d**, Proportion of differentially expressed genes in fused CDs and all genomes of BXPC3 and PANC1. P-value was calculated by fisher’s exact test. **e,** Box plots represent gene expression levels in fused CDs and outside CDs in BxPC3 and PANC1 cells. The box represents the interquartile range (IQR), with the centerline denoting the median; the whiskers extend to 1.5 times the IQR (or to the maximum/minimum if < 1.5 × IQR). Gene expression was compared as Log2FC (cancer cell line/HPDE6C7) with the P-value obtained by the Wilcoxon rank-sum test. **f**, Schematic diagram of the influence of SVs on gene expression in adjacent CDs. Left panel: The SV occurs within the CD, and the impact on gene regulation is generally restricted to this CD. Right panel: The SV occurs at a CD boundary or where CD structures are more loosely defined, and the effect on gene regulation spreads to adjacent CDs.

We then analyzed the correlation between CD fusion and differential gene expression in two cancer cell lines. The results showed that the proportion of differentially expressed genes in fused CDs was significantly higher than that in other regions of the genome (Fig. 4d, Supplementary Tables 19 and 20). More importantly, we further analyzed the interaction frequency of CDs adjacent to the cross-CDB deletion and found that the interaction frequencies of fused CDs on either side of deletion were significantly higher than those of regions outside the fused CDs (Fig. 4e), suggesting that the CD, as an essential functional unit of 3D chromatin, is able to confine the influences of SVs on 3D chromatin organization and gene expression to the adjacent CDs to maintain the structural and functional stability of the whole genome (Fig. 4f).

Collectively, these data show that cancer-specific SVs may regulate gene expression by remodeling CDs in PDAC. Moreover, the bulk remodeling effect observed in the 3D genome partly depends on intercellular genomic heterogeneity, which further expands our understanding of the pathogenesis of SVs in PDAC.

### Homozygous deletion of *CDKN2A* is associated with MIR31HG upregulation in part through concomitant amplification and contact domain fusion of adjacent genomic region

Previous studies have shown that inactivation of the tumor suppressor gene *CDKN2A* plays an important role in the occurrence and development of PDAC. *CDKN2A* inactivation occurs in approximately 90% of PDACs through various mechanisms, among which homozygous deletion is one of the most common pathways^2, 52, 53^. Our above findings confirmed the homozygous deletion of *CDKN2A* in both BxPC3 and PANC1 (Fig. 1g). Next, we explored the effect of this homozygous deletion on 3D genome organization and gene expression in chromosome 9. In view of the differences in deletion length between the two cancer cell lines, we first carried out the analysis in PANC1 and found that the interaction between adjacent CDs on both sides of the deletion was significantly enhanced to form a fused CD. Moreover, the internal interaction was also significantly intensified between adjacent CD regions, which was consistent with the finding of duplications on both sides of this deletion in the genomes by TGS and NGS (Fig. 5a and b). We further analyzed the gene expression changes in this region and found that the expression of *CDKN2A*, *CDKN2B* and *MTAP* (genes in the deletion region) had almost disappeared, whereas the expression of *MIR31HG* and *LINC01239* (genes on either side of the deletion) had been significantly upregulated (Supplementary Fig. 8a and Supplementary Table 21). However, there were no significant changes in the expression of IFNA family members, which may be attributed to the lack of transcriptional activity of alpha interferon in both cell lines without further stimulation by viral infection. Interestingly, we also found the MTAP-DMRTA1 gene fusion, which was consistent with the previous NGS results of PANC1 in the CBioPortal database. Both *MIR31HG* and *LINC01239* are long non-coding RNAs, and data from TCGA and GTEx revealed that their expression was significantly higher in PDAC tissues than in adjacent normal tissues (Fig. 5c). Previous studies have shown that *MIR31HG* presents a carcinogenic phenotype in various solid tumors, such as PDAC, squamous cell carcinoma of the head and neck, and esophageal cancer^54–56^, while the effects of *LINC01239* have rarely been reported. As *MIR31HG* is located in the fused CDs, it can be speculated that the upregulation of *MIR31HG* in PANC1 might be related to the *CDKN2A*, *CDKN2B* and partial *MTAP* deletion in combination with the amplification of adjacent genome regions on both sides. Next, we analyzed the correlation between *MIR31HG* expression and the deletion of these three genes (*CDKN2A*, *CDKN2B* and *MTAP*) in 807 cancer cell lines from the CCLE database. Although *MIR31HG* expression was not significantly upregulated in cells with the three-gene deletion and *MIR31HG* diploidy or amplification compared with cells in which *MIR31HG* and all three genes were diploid, an upward trend of *MIR31HG* expression could still be observed in the amplification group (Supplementary Fig. 8b, c and Supplementary Table 22); the lack of statistical significance may be related to the small sample sizes of the two groups. Therefore, we further analyzed 8359 pan-cancer samples from TCGA database and found that *MIR31HG* expression under different *MIR31HG* mutation status in cancer tissues with deep deletion of the three genes (*CDKN2A*, *CDKN2B* and *MTAP*) was significantly different from that in cancer tissues in which the all *MIR31HG* and three genes were diploid. Notably, *MIR31HG* expression was significantly increased in *MIR31HG* diploid and amplified cancer tissues, indicating that the upregulation of *MIR31HG* expression was significantly correlated with *MIR31HG* copy number amplification and *CDKN2A-CDKN2B-MTAP* deletion in a pan-cancer sample (Fig. 5d, Supplementary Fig. 8D and Supplementary Table 23). In addition, we also analyzed the impact of *MIR31HG* expression on the prognosis of patients with PDAC and found that the survival of patients with high *MIR31HG* expression was significantly shortened (Fig. 5e), consistent with the carcinogenic role of *MIR31HG* in PDAC.

**Fig. 5.**
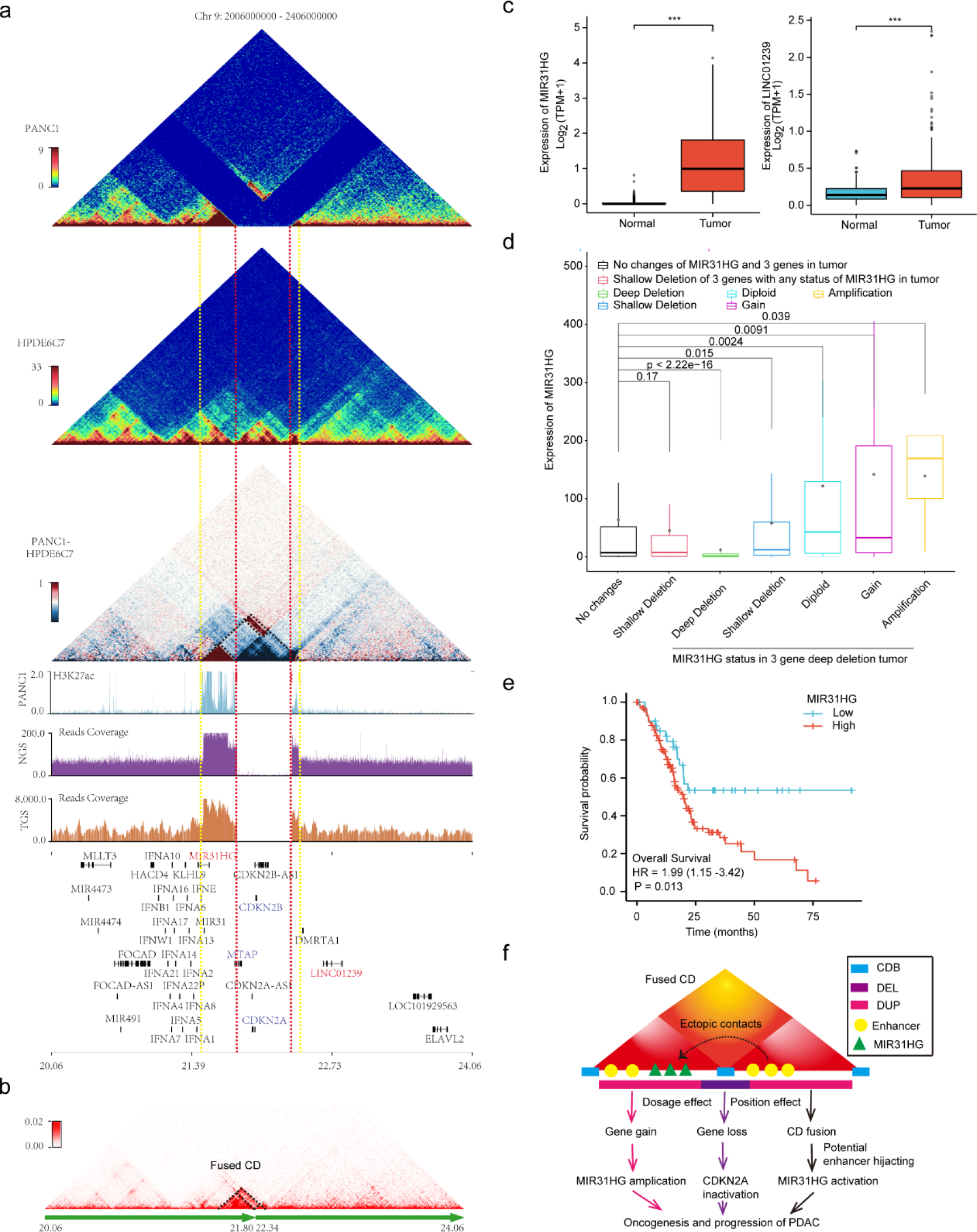
CDKN2A homozygous deletion is associated with MIR31HG upregulation partly through concomitant adjacent genome amplification and CD fusion. **a**, Diagram showing the impacts of CDKN2A homozygous deletion and concomitant amplification on 3D chromatin folding domains in PANC1. Triangle heatmaps represent chromatin contact frequency, with the top showing PANC1, middle showing HPDE6C7, and bottom showing the subtractive results. The histograms below represent the roadmap epigenome enhancer activity, marked by H3K27ac, in PANC1 (blue at top) and read coverage of next-generation (purple at middle) and third-generation (brown at bottom) sequencing for PANC1 in the same genomic region. The red dashed line denotes the break points of the homozygous deletion, and the yellow dashed line marks the boundaries of the fused CD. The black dashed line in the bottom triangle heatmap indicates the enhanced internal and external interaction of the adjacent CD. b, Triangle heatmap showing the fused CDs with the black dashed line indicated. c, Expression of MIR31HG and Linc01239 in pancreatic cancer and normal control tissues from TCGA and GTEx (n = 350). P-values were obtained by Wilcoxon rank-sum test. ***p ≤ 0.001. d, MIR31HG expression levels under different mutation states of MIR31HG and three genes (CDKN2A, CDKN2B and MTAP) in pancancer tissues from TCGA. P-values were obtained by Kruskal–Wallis test. e, Kaplan–Meier survival curves for overall survival according to MIR31HG expression in the TCGA pancreatic cancer dataset with a total of 178 cases (low group: 43, high group: 135). The P value was obtained by Cox regression in R (version 3.6.3). f, Schematic diagram showing that CDKN2A homozygous deletion could promote oncogenesis and the progression of PDAC by upregulating MIR31HG through concomitant amplification (dosage effect) and CD fusion (position effect). Dosage effects include oncogene MIR31HG amplification and suppressor CDKN2A inactivation. Position effects refer to potential enhancer hijacking through CD fusion.

Similarly, we studied the homozygous deletion related to *CDKN2A* in BxPC3. As the length of deletion was larger than that in PANC1, the expression of the *CDKNA2A*, *CDKN2B*, *MTAP* and *MIR31HG* genes, which were within the range of the deletion, was lost. At the same time, the interaction between adjacent CDs on both sides of this deletion was significantly enhanced, forming a CD fusion. However, due to the lack of expressed genes in the fused CD region, no changes in expression were observed (Supplementary Fig. 8a and e).

These findings suggested that *CDKN2A* homozygous deletion was associated with upregulation of *MIR31HG* expression in PDAC, which may be related to concomitant amplification and CD fusion in the adjacent genomic regions of *CDKN2A* homozygous deletion (Fig. 5f). In conclusion, our research revealed the effects of *CDKN2A* homozygous deletion on 3D genome organization and gene expression, providing new insight for understanding *CDKN2A* inactivation to drive the occurrence and development of PDAC.

### Identification of *SMAD4* deletion-associated complex genomic rearrangements in PDAC

*SMAD4*, one key driver gene of PDAC, is known to be lost in approximately 55% of pancreatic cancers, with homozygous deletion accounting for approximately 30% of these cases^57^. However, little is known about the effect of homozygous SMAD4 deletion on 3D genome organization. The SMAD4 homozygous deletion in BxPC3 was validated by both the TGS technique and the Hi-C method (Fig. 6a and Supplementary Fig. 2d). According to our abovementioned findings, cross-CDB deletion enhanced adjacent CD interactions on both sides of the breakpoints (Model Fig. 6b-top). Surprisingly, the interaction between the two sides of the cross-CDB deletion involving the SMAD4 gene was not enhanced but disappeared, and only a single bin on both sides of the deletion was found to have enhanced interaction at 40-kb resolution (Fig. 6a, Model Fig. 6b-bottom). To further explore the ground truth underlying these abnormal changes, we first checked the interaction heatmap of chromosome 18 of BxPC3 and found a number of enhanced distal ectopic interactions. These unusual long-range regions were mainly related to three large deletion sites, including the region of SMAD4 deletion (Fig. 6c). Next, we identified the three large deletion sites from 40 kb Hi-C matrices and divided chromosome 18 into scaffolds (Fig. 6c and d). Then, we rearranged the scaffolds to map them to the new chromosome 18 according to de novo genome assemblies based on chromatin interactions^58^. The aberrant long-range chromosomal interactions all but disappeared after the rearrangement. These results indicate that homozygous loss of *SMAD4* and multiple deletions on chromosome 18 in BxPC3 lead to huge, complex chromosomal rearrangements, including inversions and translocations. This is consistent with previous reports of chromothripsis in the same regions in approximately 11% of pancreatic cancer cases^4^ and may be the main cause of the paradoxical changes in local chromatin interactions. Interestingly, the chromothripsis and reorganization of chromosome 18 in BxPC3 seems to be homozygous. This may indicate that there exists some special preference mechanism for chromosome reconnection.

**Fig. 6.**
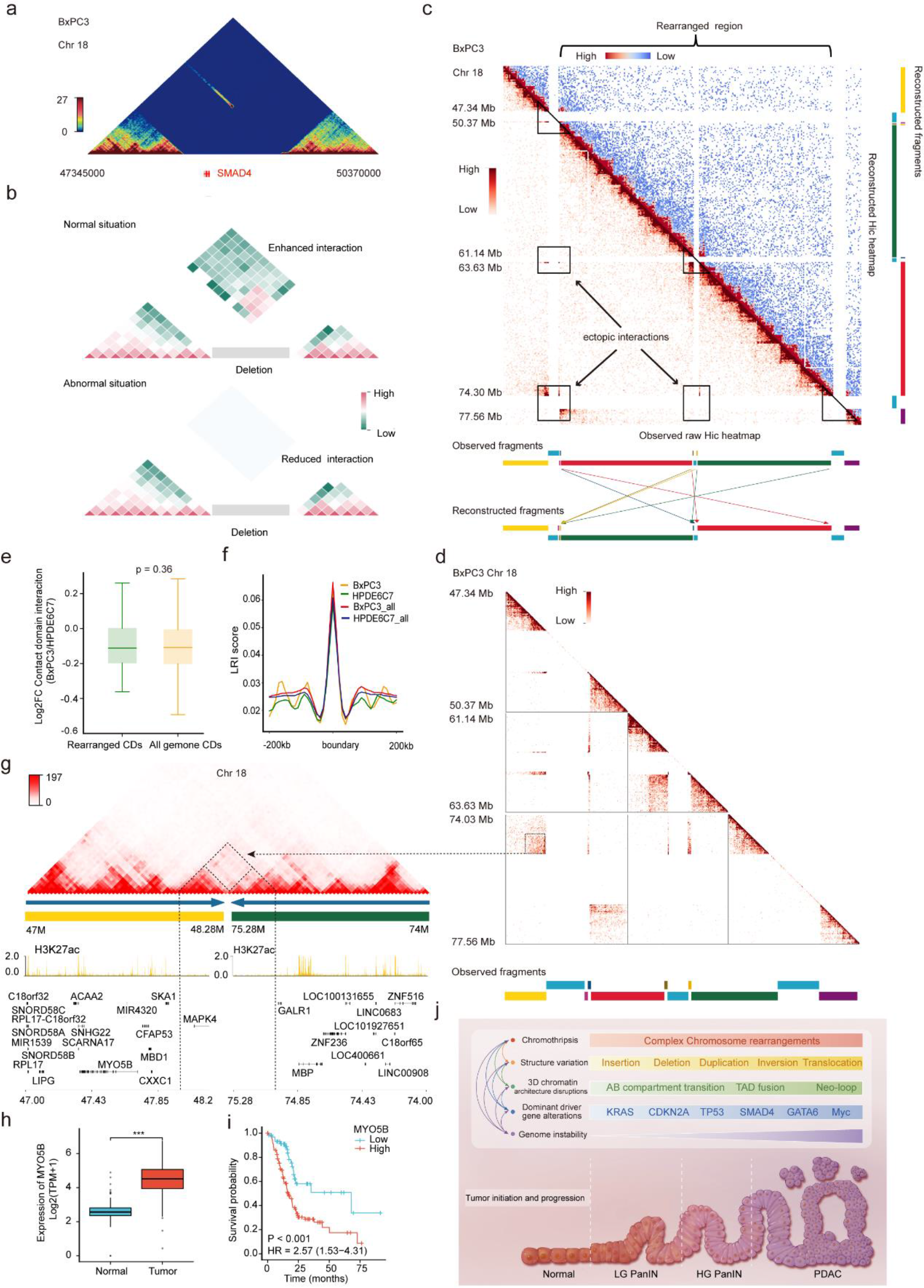
Identification of complex genomic rearrangements associated with SMAD4 deletion in PDAC. **a**, Triangle heatmap showing the interaction on both sides of the cross-CDB deletion involving SMAD4 in BxPC3. No interaction was observed on adjacent CDs of this homozygous deletion except for a single bin at 40-kb resolution. **b**, Schematic diagrams showing the impacts of homozygous cross-CDB deletion on the interaction of adjacent CDs. Top: The interaction on both sides of the cross-CDB deletion is enhanced in most normal cases. Bottom: The interaction on both sides of the cross-CDB deletion is reduced in some abnormal situations. **c**, Observed interaction heatmaps (lower-left triangle, interaction reads mapping to hg19 reference genome) and reconstructed heatmap (upper-right triangle) of chromosome 18 in the BxPC3 cell line. SMAD4 is involved in the first region marked with a triangle on the left. The aberrant chromosomal long-range interactions (indicated by black rectangles) all but disappeared after the rearrangement. Segments with different colors represent genome fragments likely resulting from chromothripsis. Arrows in different colors show simplified rearranged models of relevant segments with the same color. Briefly, red and green segments were translocated and inverted, and small segments beside them were also translocated after reconstruction. **d**, Amplification of regional heatmaps indicated by triangles and rectangles in **c**. The dashed box shows the aberrant enhanced interaction on the junction region of rearranged genome fragments corresponding to new contact domain (neo-CD) in **g**. **e**, Box plots representing the comparison of interaction frequency between CDs in the rearranged region and in the whole genome. Interaction frequency was reported as Log2FC (BxPC3/HPDE6C7). P-values were obtained by Wilcoxon rank-sum test. **f**, Comparison of the local relative insulation (LRI) score of the CDB in the rearranged region (yellow: BxPC3, green: HPDE6C7) and all genomes (red: BxPC3, blue: HPDE6C7). The LRI score was obtained by the HiCDB method. No significant difference was observed between the rearranged region and the whole genome in BxPC3 compared with HPDE6C7. **g**, Example of neo-CD formation in the junction region of rearranged yellow and green genome fragments in chromosome 18 of BxPC3 by NeoLoopFinder. The dashed triangle denotes the neo- CD corresponding to ectopic interactions in **d** with the dashed black arrow indicated. Histograms below represent roadmap epigenome enhancer activity, marked by H3K27ac, in BxPC3 (yellow). **h**, Expression of MYO5B in pancreatic cancer and normal control tissues from TCGA and GTEx (n = 350). P-values were obtained by Wilcoxon rank-sum test. ***p ≤ 0.001. **i**, Kaplan–Meier survival curves for overall survival according to MYO5B expression in the TCGA pancreatic cancer dataset with a total of 178 cases (low group: 62, high group: 116). P values were obtained by Cox regression in R (version 3.6.3). **j**, Schematic diagram showing that multiple molecular biological events are involved in the carcinogenesis and progression of pancreatic cancer. These events are not independent from each other but rather engage in crosstalk and generate a complex regulatory network.

Then, we found that the interaction frequencies of CDs in the chromosomal 18 rearrangement region did not change significantly and that the CDB remained basically unchanged (Fig. 6e and f), which is consistent with the conserved nature of CDs in the absence of one-dimensional sequence changes. However, a neo-CD was identified in the junction region of the rearranged genome fragments by NeoLoopFinder^59^ (Fig. 6g and Supplementary Fig. 9a). Subsequently, we analyzed the expression of the *MAPK4*, *DCC* and *CTDP1* genes within the neo-CD range and found that the transcription of all three genes was extremely low and did not exhibit any significant changes (Supplementary Table 24). This may be related to the fact that enhancers marked by H3K27ac within the neo-CD region had no significant alterations in their activities (Fig. 6g and Supplementary Fig. 9a). The above findings suggest that CD can reduce the disruption of 3D genomic organization caused by complex chromosomal rearrangements, maintaining the basic architecture of the chromatin and stabilizing the expression of genes within the CDs. This may be an intermediate protective mechanism by which chromatin can limit the disruption of gene expression by SVs and could be the result of adaptive selection under natural stresses during evolution.

Indeed, rather than being restricted to chromosomal rearrangement junction regions, extensive ectopic chromosomal rearrangements may also affect long-range gene regulation throughout chromosome 18 by altering the spatial location of cis-regulatory elements. We therefore screened the entire set of differentially expressed genes across chromosome 18. To identify differential gene expression related to chromosome rearrangements, we excluded those genes with no significant difference in expression or low self-expression level (FPKM < 1) in BxPC3 and PANC1. Finally, a total of 51 candidate rearrangement-related differentially expressed genes were identified, of which 39 showed higher expression in pancreatic cancer than in adjacent normal tissues and 28 were significantly related to poor prognosis (Supplementary Table 25). Notably, both *MYO5B* and *VPS4B* within the rearrangement region were upregulated in pancreatic cancer and associated with poor prognosis (Fig. 6h and I, and Supplementary Fig. 9c and d). Previous studies revealed that both of these genes encoded proteins that are involved in crucial cell biological processes such as plasma membrane recycling^60, 61^ and intracellular protein trafficking^62, 63^. Unfortunately, there have been few studies of these genes in tumors at present. Similarly, *DSC2*, *DSG2*, and *LAMA3* were also found to be highly expressed and significantly associated with poor prognosis in pancreatic cancer (Supplementary Fig. 9c and d). It has been shown that their encoded proteins are involved in epithelial cell-cell junctions, adhesion and cell motility and migration and therefore may play an important role in the carcinogenesis and progression of pancreatic cancer^25, 64, 65^.

Collectively, our study identified complex chromosomal rearrangements associated with *SMAD4* deletion on chromosome 18 and revealed their effects on 3D genome organization and related gene expression. Moreover, our data further verified the critical role of CDs in maintaining chromatin structural stability. These results provide more clues for understanding the complicated regulatory roles of SVs, 3D chromatin architecture and gene expression changes in the carcinogenesis and progression of pancreatic cancer (Fig. 6j).

## Discussion

Recently, with the rapid development and application of three-generation sequencing (TGS) and high-throughput chromatin conformation capture techniques (Hi-C), increasing evidence has shown that SVs and the 3D genome play critical roles in tumorigenesis and development^11, 12, 24, 26–28^. However, the spectrum of SVs and overall 3D genome architecture, as well as their dynamic interplay, during malignant transformation of normal pancreatic ductal epithelium remain largely undefined. In this study, we integrated multiomics data and applied TGS, in situ Hi-C, and RNA-seq technologies in combination with data from multiple public databases to perform comprehensive analyses on two PDAC cell lines and a normal immortalized human pancreatic ductal epithelial cell line. This study revealed the signatures of SVs and multidimensional alterations in the spatial organization of chromosomes in PDAC cells and further characterized the complicated interplay of 3D chromosome organization and SVs and their impacts on gene expression. These results could expand our understanding of the complex regulatory network of molecular biology involved in the carcinogenesis and progression of PDAC.

To systematically study SVs in PDAC, we first employed SMRT technology to establish the signatures of SVs in cancer cell lines and identified more than 20000 SVs. These results fully reflect the substantial advantages of TGS technologies for SV detection and identification^16^. Interestingly, up to 23035 SVs were detected in the immortalized ductal epithelial cell line HPDE6C7, slightly more than the number detected in the cancer cell lines PANC1 and BxPC3. This difference might be related to the process of immortalization of human pancreatic ductal epithelial cells derived from normal adult human pancreatic ducts transfected by the E6E7 gene of human papilloma virus^66, 67^. In addition, multiple recent studies using TGS have identified more than 20000 SVs in different normal human genomes^68–70^, indicating that SVs are polymorphic in the human genome. Such polymorphic SVs can generate novel genomic rearrangements and contribute to the maintenance of genomic diversity^20^. Moreover, most of the polymorphic SVs prevalent in the population are not pathogenic^71^. To differentiate polymorphic from potentially disease-causing SVs, it is essential to determine whether an SV occurred de novo or was inherited, as de novo (i.e., disease-specific) SVs are more frequently associated with the etiology of a disease^71^. Therefore, identifying pathogenic SVs will be of great significance and deserve more attention in future studies.

Notably, SVs also have a certain preference in their genomic distribution. First, at the linear genomic level (1D), we found that SVs were mainly distributed in intergenic and intronic regions and less distributed in exonic regions, indicating that most SVs did not alter coding sequences directly, which in turn maintains the evolutionary stability of the human genome. Second, at the 3D genome level, SVs tended to be enriched in compartment A and at the boundaries of CDs, whereas SVs in compartment B and inside CDs were relatively rare, suggesting that SV occurrence and formation may be influenced by 3D genomic organization, while SVs may exert a pathogenic effect by remodeling the genome architecture^20, 47^. Consequently, in addition to changing the gene dosage in linear genomic exonic regions^72^, SV primarily produces pathogenic effects by altering the spatial organization of chromatin to interfere with the positioning and/or copy number of regulatory elements, that is, exerting position effects^20^. Through these position effects, SV may affect the expression of genes that are distant from SV breakpoints and participate in carcinogenesis and progression. For example, deletions and duplications can not only alter the dosage of cis-regulatory elements but also affect their spatial positional distribution, which in turn may affect gene expression through higher-order chromatin organization of the locus. Similarly, inversions and translocations may affect gene expression and effect the pathogenic potential of SVs by disrupting the native enhancer regions and CDs or creating novel ones^20^, in addition to by disrupting coding sequences or producing fusion transcripts.

Previous studies have shown that SVs on the cross-CDB remove the isolation effect of the original CTCF-related boundary elements, trigger the relocation of enhancers, and may affect enhancer-promoter communication, leading to aberrant gene expression^42, 48, 49^. Deletion and duplication, as the most common simple SVs, have been shown to induce the fusion of adjacent CDs and neo-CD formation, respectively, in numerous tumors^28, 29, 46, 49^. Although similar phenomena were observed in our results, the actual situation was far more complicated than expected. We observed partial cross- CDB SVs that did not result in corresponding CD fusions or neo-CD formation. We speculate that this may be related to SV frequency or copy number variations in different cell lines, as Hi-C data are derived from the average value of a specific cell population. Fused CDs or neo-CDs within the entire cell population might be masked to varying degrees if the chromosomes of heterogeneous cancer cells do not undergo deletion or duplication of CTCF-associated boundary insulators. Accordingly, our understanding of the true nature of chromatin domains is probably obscured by intercellular genomic heterogeneity in population-averaged data. The recent development of single-cell Hi-C is exciting, as this technology is expected to solve this issue^73, 74^. More importantly, our study revealed that the structural disturbance in CDs caused by cross CDB deletions appeared to be confined to adjacent CDs, indicating that CDs tended to restrict the impact of SVs to the greatest extent and thereby maintained the stability of the overall 3D chromatin architecture. This is consistent with the evolutionarily conserved features of CDs found in the study, which might be the result of adaptive selection under natural evolutionary stress. Therefore, cross-CDB SVs may affect gene regulation by disrupting the 3D structure of adjacent CDs. However, the subsequent self-protection mechanism limits the further influence of SVs, indicating that the process of pancreatic carcinogenesis and development involves the dynamic interplay and complex regulation of multiple mechanisms.

Previous studies have identified the loss of *CDKN2A* on chromosome 9 and *SMAD4* on chromosome 18 as important drivers of pancreatic carcinogenesis and progression, with homozygous deletion being a major cause of inactivation of these two key tumor suppressor genes^75^. In this study, we first confirmed the presence of *CDKN2A*- and *SMAD4*-related homozygous deletions in PDAC cells using TGS. Then, through Hi-C sequencing, we found that the homozygous deletion was accompanied by ectopic chromosome rearrangement. Furthermore, we revealed the effects of this complex chromosome rearrangement on 3D genome organization and gene expression alterations, providing new biological insights and potential therapeutic possibilities for understanding the carcinogenesis and progression of PDAC. First, we identified a homozygous *CDKN2A-CDKN2B-MTAP* codeletion along with tandem duplications of the flanking regions on chromosome 9 in the PANC1 cell line. This complex SV led directly to the loss of the *CDKN2A* tumor suppressor signaling pathway and significantly upregulated the expression of the *MIR31HG* gene in its neighboring region. This could be the result of cross CDB deletion-related adjacent CD fusion and enhancer hijacking^49, 76, 77^, as well as of an increase in gene dosage at the 1D level caused by tandem duplications within CDs. In practice, deletions and tandem duplications of specific cancer-related genes are also quite common in other tumors^75^. These results indicate that SV can not only alter gene dosage at the 1D level but also regulate the expression of cancer-related genes through 3D position effects, which collectively contribute to pancreatic carcinogenesis and progression. Previous studies have shown that MIR31HG can promote cancer in a variety of solid tumors^54–56^. Therefore, the development of targeted therapeutic strategies may have good prospects in clinical applications.

Our study also showed that the deletion of a region on chromosome 18 of BxPC3 cells produced an unexpected result on Hi-C; *i.e*., the interaction between the regions flanking the lost *SMAD4* gene was not enhanced but basically disappeared, which could not be explained by simple deletion. Further localized scaffold rearrangements on chromosome 18 confirmed the complex ectopic rearrangements of the genome, including the *SMAD4* loss region. Recently, an increasing number of studies have shown that extensive chromosomal rearrangement caused by complex SV is a critical mechanism involved in the genetic instability of PDAC^4, 78, 79^. These findings challenge the current model of PDAC tumorigenesis and provide novel insights into the mutational processes that give rise to these aggressive tumors. However, these studies have focused on the alterations of 1D chromosome structure and their effects on the regulation of gene expression. Based on this knowledge, we further analyzed the chromatin interactions in the rearranged regions of chromosome 18 and found that the interaction frequencies did not change significantly within the rearranged regions that did not involve the CD boundary, suggesting that the presence of an isolated boundary may be crucial for maintaining the stability of CD organization. This is because genomic rearrangements that do not involve boundaries are more likely to alter gene dosage within CDs^20^, not the ectopic contacts between adjacent CDs. However, enhanced interactions and neo-CDs were observed in the junction loci of the chromosomal rearranged segments, but the expression of the MAPK4, DCC, and CTDP1 genes within the neo-CD region did not change dramatically, indicating that neo-CD formation is not always associated with gene expression changes, especially when *cis* regulatory elements remain stable^59, 80^. Accordingly, gene regulation in PDAC genomes is sophisticated and influenced by multiple factors. Disruption of chromatin folding domains caused by chromosome rearrangement may contribute to gene expression changes, but this is not always the case^81^.

Next, we analyzed the differentially expressed genes associated with genomic rearrangements across the entirety of chromosome 18 and found that the vast majority of them (39/51) were upregulated in PDAC and were significantly related to poor prognosis. DSC2 and DSG2 are typical examples. As an important component of the desmosome and the most widely distributed isoform of desmocollin (DSC), desmocollin 2 (DSC2) has been demonstrated to be essential for the adhesion of epithelial cells and serves as a vital regulator in signaling processes such as epithelial morphogenesis, differentiation, wound healing, cell apoptosis, migration, and proliferation^82^. Recent studies have suggested that the aberrant expression or disruption of DSC2 might lead to some disorders, including heart disorders^83^, certain cancers^84–86^, and other human diseases^87^. In addition, the desmoglein 2 (DSG2) gene product is a calcium-binding transmembrane glycoprotein component of desmosomes in vertebrate epithelial cells. Previous studies have shown that DSG2 has good prognostic value in several solid tumors, such as cervical cancer ^88^, anaplastic thyroid cancer^89^, and ovarian cancer^90^. A recent study suggested that DSG2 could promote the carcinogenesis and progression of squamous cell carcinoma (SCC) by enhancing exosome synthesis and secretion^91^. In addition, laminin subunit alpha 3 (LAMA3) belongs to the laminin family of secreted molecules and is responsive to several epithelial-mesenchymal regulators, including keratinocyte growth factor, epidermal growth factor and insulin-like growth factor. The methylation or aberrant expression of LAMA3 is significantly associated with poor prognosis in patients with pancreatic and ovarian cancer^92, 93^. Consequently, these chromosomal rearrangements associated with differential gene expression may serve as prognostic markers and potential therapeutic targets for PDAC in the future. In short, complex genomic rearrangements that occur on the chromosome may affect cancer-related genes by altering the 3D genome organization and participate in the carcinogenesis and progression of PDAC. This is an important addition to previous findings. Our results provide a new and high-dimensional perspective for a more comprehensive understanding of the molecular biological processes involved in pancreatic carcinogenesis and development. However, why chromosomal rearrangements occur so frequently in PDAC remains difficult to answer; it is speculated that this may be a result of selective adaptation exerted by extensive desmoplasia.

Notably, an SV manifests as a ‘junction’ between two ‘breakpoints’ in the genome. Only when TGS sequencing reads cross and cover the two break points of the SVs can the SV be identified by TGS SV calling software. In this study, we failed to identify another two large deletions related to chromosome rearrangements not involving SMAD4 using this software. However, this information could be easily obtained by using a Hi-C heatmap, suggesting that the current TGS algorithms have limitations in identifying complex SVs associated with chromosome rearrangement. Therefore, the combined application of TGS SV calls, Hi-C interactive data and even bionano data and/or genome assembly algorithms are necessary approaches and effective strategies to accurately interpret complex genomic SVs for PDAC in the future^69^.

In summary, our research applied multiomics techniques to establish the signatures of SVs and 3D genome architecture and characterize the dynamic interplay between them in PDAC. Furthermore, the impact of homozygous deletion of two key driver genes, CDKN2A and SMAD4, on 3D chromatin folding domains and the expression of related genes in the carcinogenesis and progression of PDAC were specifically elucidated. These findings provide a new spatial perspective toward a comprehensive understanding of the functions and pathogenic mechanisms of SVs in pancreatic carcinogenesis and development. However, it must be acknowledged that there are also some limitations in this study, such as the representativeness of the cell lines and the limitations of TGS technology. Therefore, the conclusions of this study still need to be validated with more basic experiments and in clinical samples. In short, the current research based on high-dimensional genome will contribute to identifying new molecular markers or potential targets for PDAC and providing new precise targeted regimens, which is of great practical significance to improve the therapeutic challenges of PDAC with an extremely poor prognosis.

## Materials and Methods

### Cell lines and culture

Human pancreatic ductal epithelial cells (HPDE6-C7) and the human pancreatic cancer cell lines PANC1 and BxPC3 were obtained from the American Type Culture Collection (ATCC) (https://www.atcc.org/). All cell lines were cultured under recommended conditions, and PANC1/BxPC3 was authenticated by high-resolution small tandem repeat (STR) profiling. Transcriptome cluster analysis was performed on three cell lines in the CCLE+GSE97003 database, which matched well with the public database.

### Identification of structural variations

Genomic DNA was extracted from the cell lines BxPC3, PANC1 and HPDE6C7 using a QIAamp DNA Mini Kit/DNeasy Plant Mini Kit1 (QIAGEN). The integrity of the DNA was determined with an Agilent 4200 Bioanalyzer (Agilent Technologies, Palo Alto, California). Eight micrograms of genomic DNA was sheared using g-Tubes (Covaris) and concentrated with AMPure PB magnetic beads. Each SMRT bell library was constructed using the Pacific Biosciences SMRTbell Template Prep Kit 1.0 Express Template Prep Kit 2.0. The constructed library was size-selected by the Sage ELF BluePippin™ system for molecules 8-12 kb and 14-17 kb, followed by primer annealing and the binding of the SMRT bell templates to polymerases with the DNA Polymerase Binding Kit. Sequencing was carried out for 30 h on the Pacific Bioscience Sequel II platform by Annoroad Genomics. We used NGMLR (https://github.com/philres/ngmlr) to perform the alignment with default parameters. We used sniffles^16^ to determine structural variation by using default parameters and identified cancer-specific structural variation by BEDTools^94^. The SV classification algorithm is comprehensively defined in another study^101^. Cancer genomes are shaped with both simple SVs and complex SVs. In this work, the complex SVs are defined as local assemblies that are made up of multiple SV junctions from different genomic locations on the reference genome, while the simple SV assemblies are defined as assemblies that only contain single junction event.

### In situ Hi-C

#### Hi-C library preparation and sequencing

We used 40 ml 2% formaldehyde solution to crosslink the seed tissue for 15 min at room temperature in vacuum. Then, we added 4.324 ml of 2.5 M Gly to quench the cross-linking reaction. The supernatant and tissues were removed from the precipitate. We kept the tissue in liquid nitrogen. Tissues were resuspended in 25 ml extraction buffer (10 mM Tris-HCl at pH 8, 10 mM MgCl2, 0.4 M sucrose, 0.1 mM phenylmethylsulfonyl fluoride [PMSF], 5 mM b-mercaptoethanol, and 13 protease inhibitors; Roche). Next, the suspension was filtered through Miracloth (Calbiochem). We then performed centrifugation at 4000 rpm at 4°C for 20 min. After that, the pellet was resuspended in 1 ml extraction II buffer (10 mM Tris-HCl pH 8, 1% Triton X-100, 0.1 mM PMSF, 0.25 M sucrose, 10 mM MgCl2, 5 mM mercaptoethanol, and 13 protease inhibitors). Later, the solution was centrifuged for 10 min at 14000 rpm at 4°C. The pellet was resuspended in 300 μl extraction buffer III (10 mM Tris-HCl pH 8, 1.7 M sucrose, 2 mM MgCl2, 1 μL protease inhibitor, 0.1 mM PMSF, 5 mM b-mercaptoethanol, 0.15% Triton X-100). Afterward, another 300 μl of clean extraction buffer III was loaded and centrifuged at 14000 rpm for 10 min. The supernatant was removed, and the pellet was washed with 500 µl ice-cold 1x CutSmart buffer and centrifuged at 2500 g for 5 min each time. The remaining pellets contained the nuclei. Then, we used restriction enzyme buffer to wash the pellet twice and moved it to a safe-lock tube. The next step was to solubilize the chromatin with dilute SDS and incubate it at 65°C for 10 min. Then, the SDS was quenched by Triton X-100 overnight. Next, the nuclei were digested by 4 cutter restriction enzymes (400 units MboI) at 37°C on a rocking platform. The next step was to mark the DNA ends by biotin-14-dCTP and then perform blunt-end ligation between cross-linked fragments. Thus, the ligation enzyme could ligate the proximal chromatin DNA. We incubated the nuclear complexes with proteinase K at 65°C, and then the nuclear complexes were reverse cross-linked. DNA was then purified by phenol–chloroform extraction. We used T4 DNA polymerase to remove biotin-C from nonligated fragments. Next, we used sonication to shear the fragments to 200-600 base pairs and repair the fragments with a mixture of T4 DNA polymerase, Klenow DNA polymerase and T4 polynucleotide kinase. Biotin-labeled DNA fragments were then enriched by streptavidin C1 magnetic beads. The Hi-C library from the beads was sequenced on the Illumina HiSeq X Ten platform with 150 bp paired-end reads. Raw reads were trimmed to 50 bp and then filtered by fqtools plus (https://github.com/annoroad/fqtools_plus) to discard the reads with adapters (> 5 bp adapter nucleotide) and a high N ratio (>5%) and low-quality reads.

#### Hi-C reads mapping and normalization

Clean reads were mapped to the *Homo sapiens* genome assembly (hg19) using Bowtie2 (v2.3.4)^95^. An optimized and flexible pipeline filtered out unmapped, multimapped, or invalid paired-end reads by Hi-C Pro^96^. Only uniquely valid paired-end reads were retained for analyses. The interaction matrices at various resolutions (i.e., with the genome partitioned into bins of different sizes) were constructed using HiC-Pro software (v2.7.1) with default settings^96^. Hi-C interaction matrices were constructed with bin sizes of 1 Mb, 100 kb, 40 kb, 20 kb, 10 kb, and 5 kb at the genome-wide level following the methods of HiC-Pro software (v2.7.1) with default settings^96^. Briefly, we utilized an improved computational efficiency ICE^96^ (Iterative Correction) (https://github.com/seqyuan/iced) method to remove potential Hi-C interaction bias. The genome-wide Hi-C resolution values were calculated based on the interaction maps according to previously published definitions^95^.

#### Differential interaction and calculation of chromatin interaction and IDE values

To analyze the difference between interaction matrices in the BxPC3, PANC1, and HPDE6C7 samples, the different matrices were computed by subtracting the z-score matrices of sample-paired intra-interaction matrices. To measure the distance-dependent decay in the cis-interactions of a genome, we used 1 Mb interaction matrices. The intrachromosome interaction frequency between each bin with the same distances on the reference genome was calculated as real distance in space (https://github.com/dekkerlab/cworld-dekker). Interaction frequencies were log10 transformed to fit a linear model. The slope of each model was outputted as the corresponding finalized IDE value (https://github.com/dekkerlab/cworld-dekker).

### Identification of A/B compartments

Briefly, we calculated the expected score within each matrix using loess smoothed averaging over the intrachromosomal interactions. Then, we obtained the observed/expected ratio of intrainteraction matrices. Then, we constructed a Pearson’s correlation matrix reflecting the chromosomal interactions for each pair of bins, which was used for principal component analysis (PCA). We first calculated the eigenvalues of the first and second principal components for each chromosome. Then, we found the A/B compartments according to the positive and negative eigenvalues of each bin and adjusted by gene density. After that, we checked whether the gene expression level in each type of compartment in each chromosome corresponded to the definition of A/B compartments, in which the A compartment gene expression level was higher and the B compartment gene expression level was lower. We also checked the iced matrices and Pearson’s correlation matrices of each chromosome. Finally, we determined which chromosome uses eigen2 values and which chromosome uses eigen1 values in each sample and each chromosome. In addition, we also determined whether -1 was multiplied in each chromosome’s eigenvalue. In the end, for each chromosome, the A/B compartments were determined to have higher gene expression level in A compartment and lower gene expression level in B compartment, and we were able to identify different types of compartment interaction patterns in the iced matrix and Pearson’s correlation matrix (https://github.com/dekkerlab/cworld-dekker).

### Identification of CDBs (HiCDB)

HiCDB was applied to identify CDBs. We used the matrix at 10-kb resolution as input and ran HiCDB with default parameters. After we found the boundaries, we compared the LRI score of each boundary between each sample. When one sample’s boundary region LRI score was twice that of the other samples, we considered it to be a specific boundary^43^.

### Loops

Genome-wide chromatin loops were identified using Hi-C Computational Unbiased Peak Search (HiCCUPS) as part of the Juicer package using 5-kb bins and default parameters^97^. To find loops specific to one sample, we first calculated the observed/expected ratio of the two ends of the loop for each sample. Then, we made a comparison. If one sample’s observed/expected ratio was twice that of the other sample, then we consider the loop to be specific to that sample.

### RNA-seq

Total RNA of three cell lines was extracted by the TRIzol method, and libraries were constructed according to a standard protocol (Illumina) and sequenced on the Illumina HiSeq X-ten system. Three biological replicates were conducted for each library. The resulting filtered reads were aligned against the hg19 reference genome using HISAT2^98^ with default parameters (v2.1.0), and the expression level of each gene was normalized by the method developed by Traver Hart^99^, which is based on the fragments per kilobase per million mapped fragments (FPKM) values. Genes were divided into three groups: highly expressed genes (above the mean), intermediate expressed genes (between the median and the mean), and low expressed genes (below the median). Differentially expressed genes (DEGs) were identified with the DESeq2^100^ package. Genes with a Benjamini–Hochberg adjusted q-value < 0.05 and an absolute log2-fold change ≥ 1 were considered differentially expressed.

### ChIP and data analysis

ChIP-seq reads (NCBI <u>PRJEB27863</u>)^101^ were aligned to the reference genome using bowtie2^95^ software, and only unique and nonduplicated mapped reads were used for the downstream analysis. The read coverage and depth were calculated by SAMtools^102^. To examine the reproducibility of the ChIP-seq experiments, deeptools was used to generate the correlation plot for all samples, including input samples. Signal track files in BigWig format were generated using deeptools^94^ bamCoverage function and were normalized to 1 million reads for visualization. DeepTools was also used to plot the gene body and flanking region heatmap graph using the normalized signal intensity. MACS2 was used to call peaks, followed by peak annotation using bedtools^94^. Differential analysis between treated and control samples was conducted using bedtools. Functional analysis, such as GO and KEGG, for differential peak-related genes was performed with in-house scripts.

Enrichment was assessed using deepTools2 (v3.1.2)^103^ with default parameters.

### The distribution of SVs along the 3D genome

The dynamic chromosome regions were obtained through the comparison of cancer and healthy cell lines. By comparing BXPC3 (or PANC1) to the HPDE6C7 cell line, the chromosome regions can be divided into stable A/B compartments, A-B compartments, and B-A compartments. By comparing BXPC3 (or PANC1) to the HPDE6C7 cell line, the chromosome regions can be divided into cancer-gained TAD boundaries, cancer-lost TAD boundaries, and stable TAD boundaries. Cancer-specific SVs refer to those occurring in BXPC3 (or PANC1) but not in HPDE6C7. Cancer-specific SVs and dynamic chromosome regions were identified by BEDTools. To explore the distribution of SVs, we compared the density of SVs in different chromosome regions, including A/B compartments, TADs and TAD boundaries. The density of SVs is defined as the number of SVs divided by the length of each chromosome region. SV enrichment was evaluated by comparing the proportion of SVs falling in the region of interest to that in the background, which was performed via the prop.test function in R. The background density refers to the number of SVs divided by the length of the whole genome.

### GO

We applied a Gene Ontology method for functional enrichment analysis of the biological processes of the identified genes of interest, such as genes in A/B switch areas, differential loops or CDs. For each GO term, we obtained a p-value corresponding to a single, independent test and then used the BH method to correct the p values^104^.

### Neo-CDs identification

To identify new CDs in chromosome rearrangement areas, we applied NeoLoopFinder to help us find newly emerged CDs in areas that have inversions, translocations and deletions^59^.

### Statistics and data visualization

We applied the rank sum test and Fisher’s test to determine the relationship between Hi-C, transcriptome and SV data. We used Annoroad Browser (https://github.com/Spartanzhao/Annoroad-Browser) to produce the track profiles in joint multi-omics visualization of the data.

### URLs

NCBI,https://www.ncbi.nlm.nih.gov.GEPIA2, http://gepia2.cancer-pku.cn/#general.cBioPortal,https://www.cbioportal.org.GTEX,https://www.ncbi.nlm.nih.gov.GEPIA2, http://gepia2.cancer-pku.cn/#general.cBioPortal,https://www.cbioportal.org.GTEX,https://www.gtexportal.org/home/index.html. TCGA, https://www.tcga.org. XENA, https://xena.ucsc.edu. Arrayexpress,https://www.ebi.ac.uk/arrayexpress.CCLE,https://portals.broadinstitute.org/ccle/about.GSE97003,https://www.ncbi.nlm.nih.gov/geo.ExPASyhttps://web.expasy.org/cellosaurus. IGV, https://www.igv.org. ATCC, https://genomes.atcc.org/.

### Availability of data and materials

All raw and processed sequencing data generated in this study have been submitted to the NCBI Gene Expression Omnibus (https://www.ncbi.nlm.nih.gov/geo/) under accession code GSE185069. Biological material used in this study can be obtained from authors upon request. We used Annoroad Browser (https://github.com/Spartanzhao/Annoroad-Browser) to produce the track profiles in joint multi-omics visualization of the data.

### Finding

This work was supported by CAMS Innovation Fund for Medical Sciences (CIFMS No. 2016-I2M-1-001) and National Natural Science Foundation of China (81972314 and 81802463).

## Acknowledgments

Part of the results are based on the datasets generated by the TCGA team. We thank Prof. Cheng Li from Center for Statistical Science (Peking University) and Prof. Yang Chen from MOE Key Laboratory of Bioinformatics (Tsinghua University) for their constructive advice and guidance during this project implementation. We thank Prof. Cheng Li and Dongxin Lin for valuable comments and critical review of the manuscript and American Journal Experts (AJE) for assisting in the preparation of this manuscript.

## Author contributions

Yongxing Du and Chengfeng Wang conceived, designed and supervised the study with input from Hebing Chen and Zan Yuan. Yongxing Du, Hebing Chen, Zan Yuan and Yue Zhao designed and performed most of the computational analyses with help from Xiaohao Zheng, Zongting Gu and Zongze Li. Zongting Gu, Zongze Li and Yue Zhao designed and performed most of the experiments with help from Zan Yuan and Yongxing Du. Yongxing Du, Zongting Gu and Zongze Li wrote the manuscript. All the authors discussed the results and commented on the manuscript. Xiaochen Bo critically revised the manuscript. Hebing Chen and Chengfeng Wang were working group or project leader.

## Supplementary materials

### Supplementary Figures and figure legends

**Supplementary Fig. 1.**
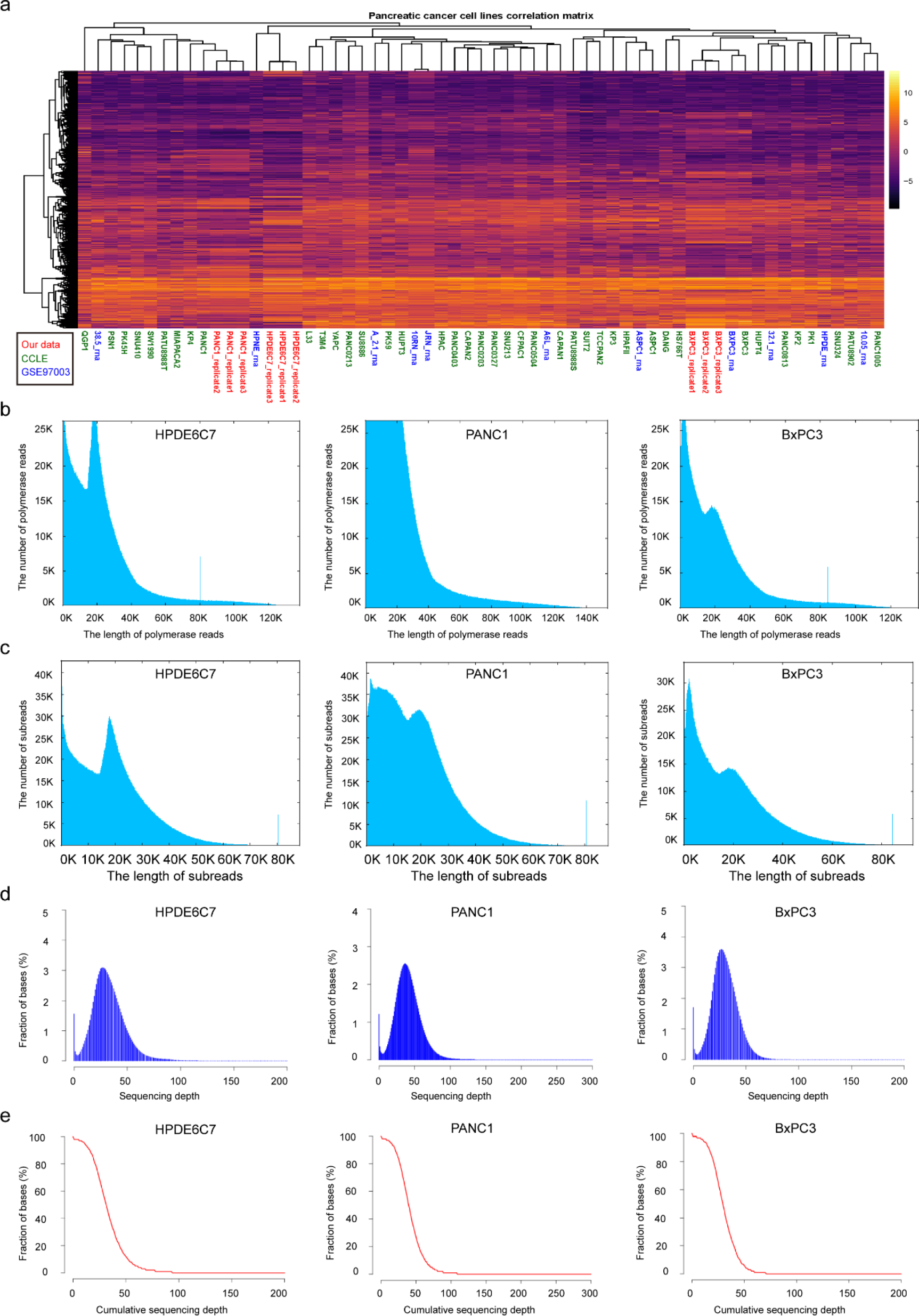
Cell line identification and quality control of long-read sequencing data. **a**, PDAC cell line correlation matrix for the clustering of our data, CCLE and GSE97003. **b**, Histogram showing polymerase read lengths of HPDE6C7, PANC1 and BxPC3. **c**, Histogram showing subread lengths of HPDE6C7, PANC1 and BxPC3. **d**, Fraction of bases as a function of sequencing depth. **e**, Fraction of bases as a function of cumulative sequencing depth.

**Supplementary Fig. 2.**
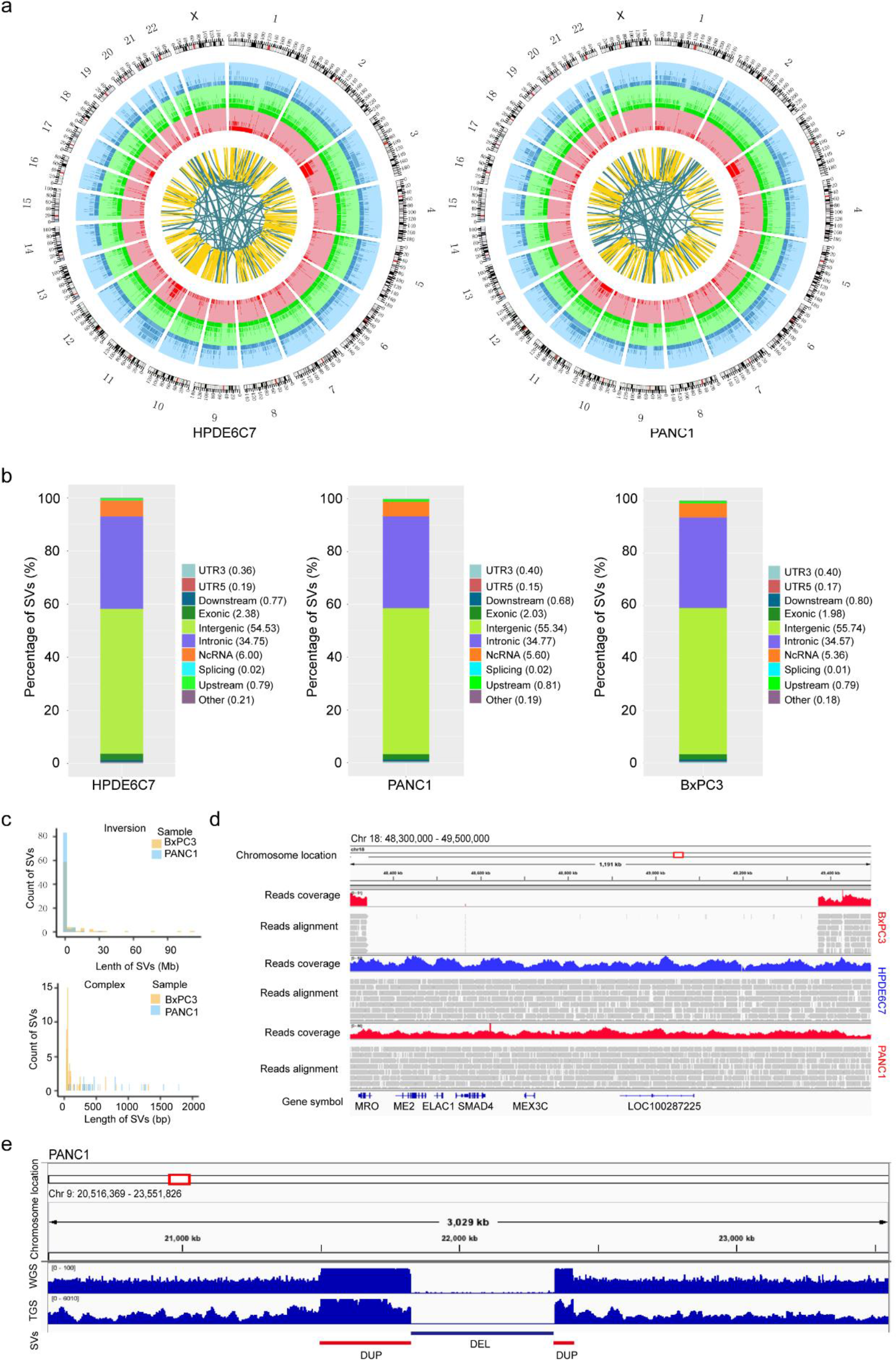
The overall landscape of SVs in PANC1, BxPC3 and HPDE6C7. **a**, Circos plots showing the high-confidence SVs detected by Sniffles in HPDE6C7 and PANC1 with 23 chromosomes inputted. The tracks from the outer to the inner circles are the chromosome coordinates, deletions, insertions, duplications and translocations. **b**, Distribution of SVs in different regions of the genome in HPDE6C7, PANC1 and BxPC3. **c**, Specific SV size histogram of PANC1 and BxPC3 found by SMRT variant calling for inversions and complex SVs. **d**, IGV image showing a homozygous deletion of SMAD4 in chromosome 18 of BxPC3. **e**, IGV image showing duplications in the region adjacent to the homozygous deletion covering CDKN2A, CDKN2B, and MTAP in PANC1.

**Supplementary Fig. 3.**
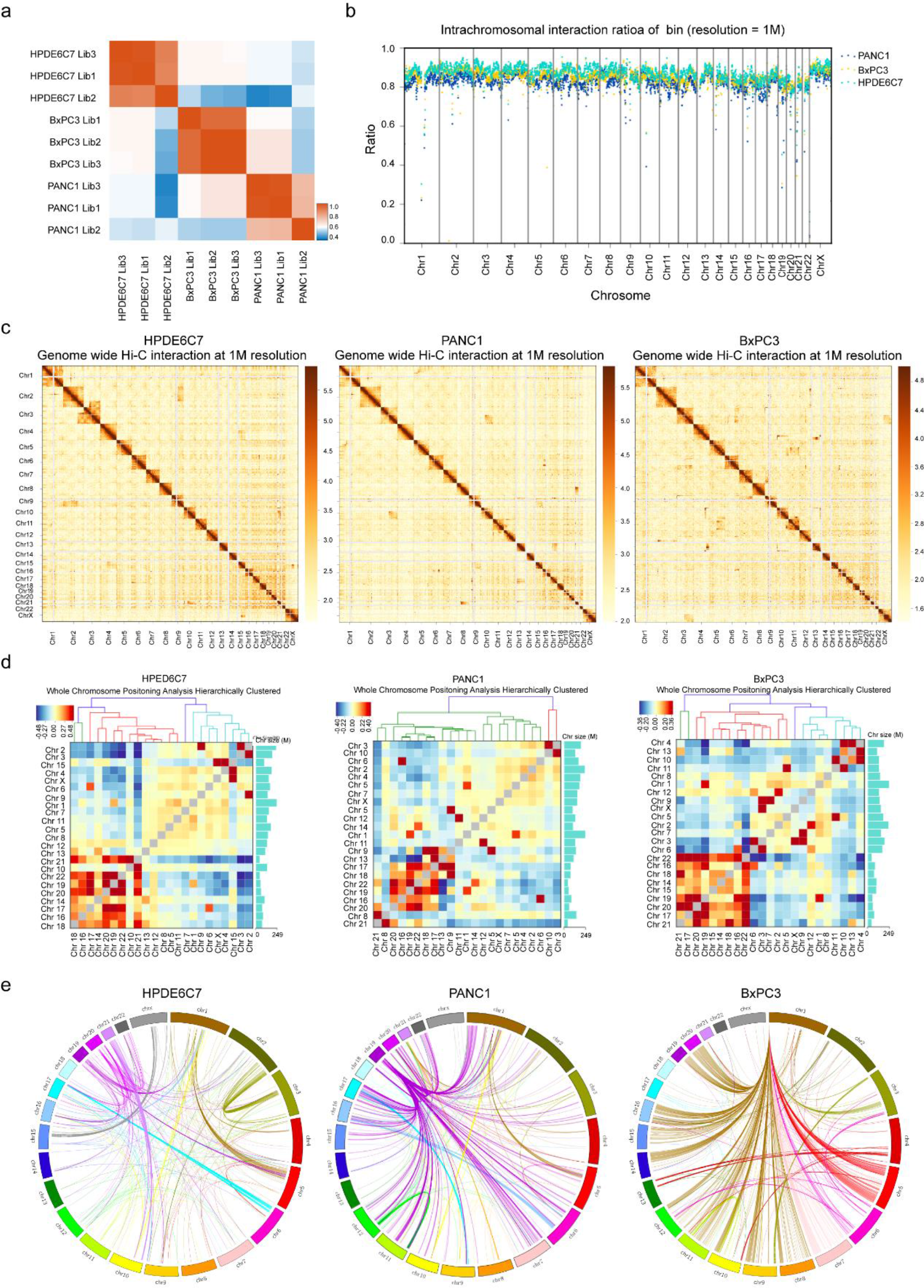
Analysis of whole chromosomal interactions in three different cell lines. **a**, Hi-C data normalization and initial quality control. Red indicates stronger interactions, and blue indicates weaker interactions. The chromosome Hi-C interaction data of the three samples from different libraries showed good repeatability. The two cancer cell types were more similar and could be distinguished from the normal HPDE6C7 cell line. **b**, Intrachromosomal interaction ratio at 1-Mb resolution in the three cell lines. BxPC3, PANC1 and HPDE6C7 exhibited relatively small differences in the intrachromosomal interaction ratio. **c**, Genome-wide all-by-all Hi-C interaction heatmap. **d**, Inter-chromosomal interactions between all pairs of chromosomes in HPDE6C7, PANC1 and BxPC3. Each block represents observed/expected interactions between chromosomes. Red and blue indicate enriched and depleted, respectively. The histogram on the right shows the chromosome size; the coordinates are in M units. **e**, Interchromosomal interaction at 40-kb resolution in each of three cell lines. The Circos plot shows the interaction of the first 1000 pairs of bins with the highest interaction strength between chromosomes. The curve indicates the position of the 1000 bin pairs with the strongest interactions.

**Supplementary Fig. 4.**
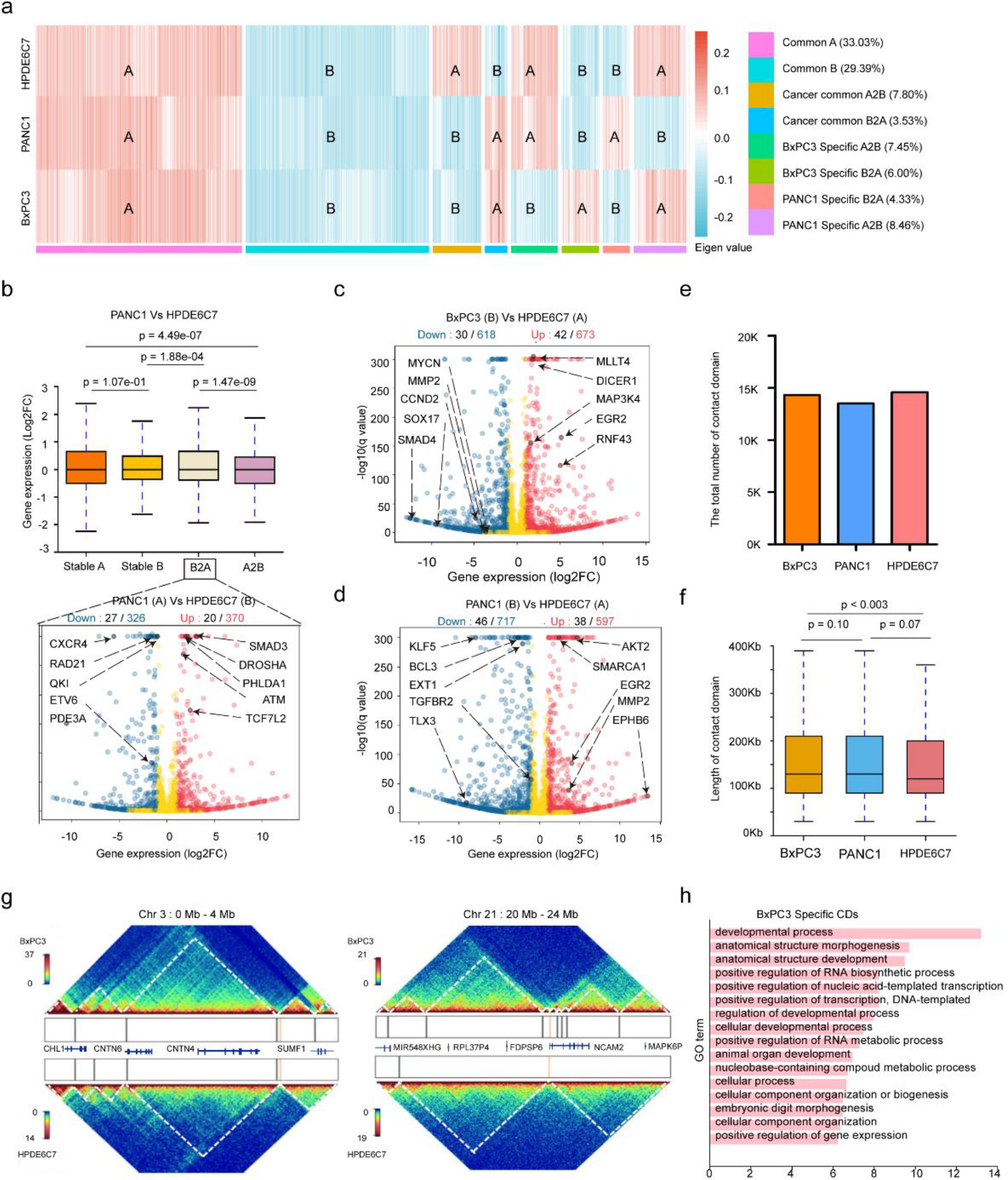
3D chromatin architecture is remodeled and correlates with gene expression changes in human PDAC. **a**, Compartment analysis using eigenvalues on all Hi-C datasets. Different categories of cancer-specific/common compartment switches were identified by comparing eigenvalues between BxPC3, PANC 1 and HPDE6C7. **b**, Top: Box plots show the gene expression comparison in different compartments between PANC1 and HPDE6C7. The box represents the interquartile range (IQR), with the centerline denoting the median, and the whiskers extend to 1.5 times the IQR (or to the maximum/minimum if < 1.5 × IQR). Bottom: Volcano plots show the number of differentially expressed genes (blue) and cancer-related genes (black) among them in the B-to-A compartment shift region. The genes indicated by the black arrow are examples of upregulated (red on the right) or downregulated (blue on the left) cancer-related genes (|Log2FC|>1 and adjusted p value<0.05). Gene expression was compared as Log2FC (BxPC3/HPDE6C7) with P-values obtained by the Wilcoxon rank-sum test. **c**, **d**, Volcano plots show the number of differentially expressed genes (blue) and cancer-related genes (black) among them in the A-to-B compartment shift region comparing BxPC3, PANC1 and HPDE6C7. The genes indicated by the black arrow are examples of upregulated (red, right) or downregulated (blue, left) cancer-related genes (|Log2FC|>1 and adjusted p value<0.05). Gene expression was compared as Log2FC (BxPC3/HPDE6C7) with the P-value obtained by the Wilcoxon rank-sum test. **e**, Histograms showing the number of CDs from HiCDB interaction matrices at 10-kb resolution in the three cell lines. **f**, Box plots represent the length of CDs from HiCDB interaction matrices at 10-kb resolution in the three cell lines. P-values were obtained by Wilcoxon rank-sum test. The CD length of the two cancer types was marginally longer than that of HPDE6C7, and there was no significant difference between the two cancer cell lines. **g**, Examples of CD alterations in regions (left chr 3: 0-4 Mb, right chr 21: 20-24 Mb) comparing BxPC3 and HPDE6C7. The vertical bars in the box between the heatmaps represent CDBs. The genes involved in the region are marked. Compared with HPDE6C7, the number and length of CDs in BXPC3 show different patterns in different chromosome regions. **h**, Biological process enrichment of differentially expressed genes located in specific CDs of BxPC3. The P values were obtained via Fisher’s exact test using Enrich R. GO, Gene Ontology.

**Supplementary Fig. 5.**
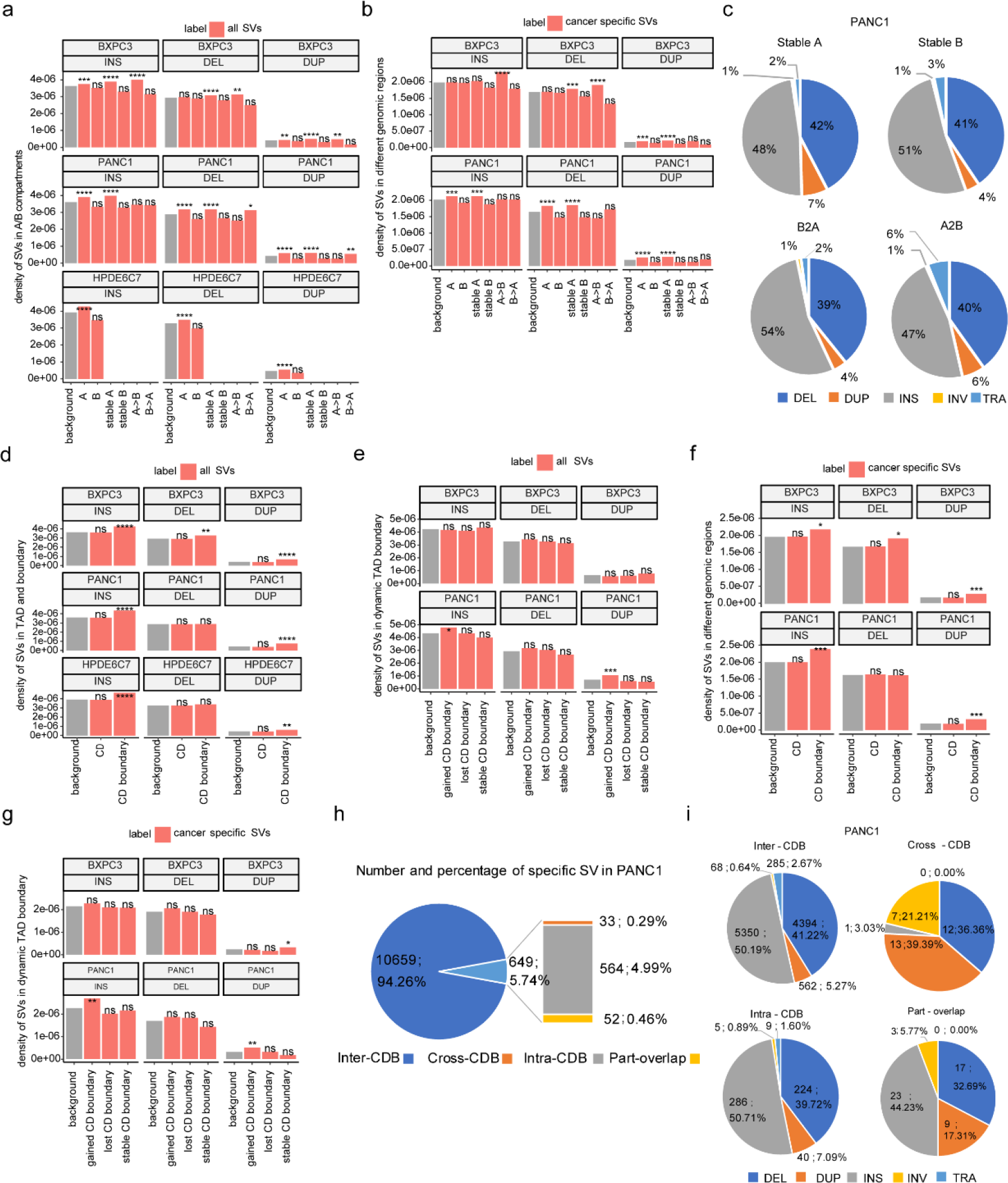
Distributions of structural variations among 3D genome architectures. **a**, **b**, The density of SVs (insertions, deletions and duplications) in A/B compartments. By comparing BxPC3 (or PANC1) to HPDE6C7, the chromosome regions can be divided into stable A, stable B, A-to-B, and B-to-A compartments. Cancer-specific SVs refer to those occurring in BxPC3 (or PANC1) but not in HPDE6C7. ****p ≤ 0.0001, ***p ≤ 0.001, **p ≤ 0.01, *p ≤ 0.05. **c**, The proportion of different types of cancer-specific SVs (less than 2 Mb length) in different A/B compartment regions of PANC1 with insertions, deletions and duplications accounting for similar percentages in stable A, stable B, A-to-B, and B-to-A compartments. **d**, **e**, The density of SVs (insertions, deletions and duplications) in CDs and CDBs. By comparing BxPC3 (or PANC1) to HPDE6C7, the chromosome regions can be divided into cancer-gained CDB, cancer-lost CDB, and stable CDB. Enrichment was evaluated by comparing the proportion of SVs falling in the region of interest and the proportion of the length of the region of interest in the whole genome. ****p ≤ 0.0001, ***p ≤ 0.001, **p ≤ 0.01, *p ≤ 0.05. **f**, **g**, The density of SVs (insertions, deletions and duplications) in dynamic CD boundaries. Enrichment was evaluated by comparing the proportion of SVs falling in the region of interest and the proportion of the length of the region of interest in the whole genome. ****p <= 0.0001, ***p ≤ 0.001, **p ≤ 0.01, *p ≤ 0.05. **h**, Number and percentage of specific SVs in PANC1. **i**, The proportion of different types of four categories of cancer-specific SVs in PANC1according to Fig. 3d. The composition of cancer-specific SVs was roughly similar except for the Cross-CDB SV group.

**Supplementary Fig. 6.**
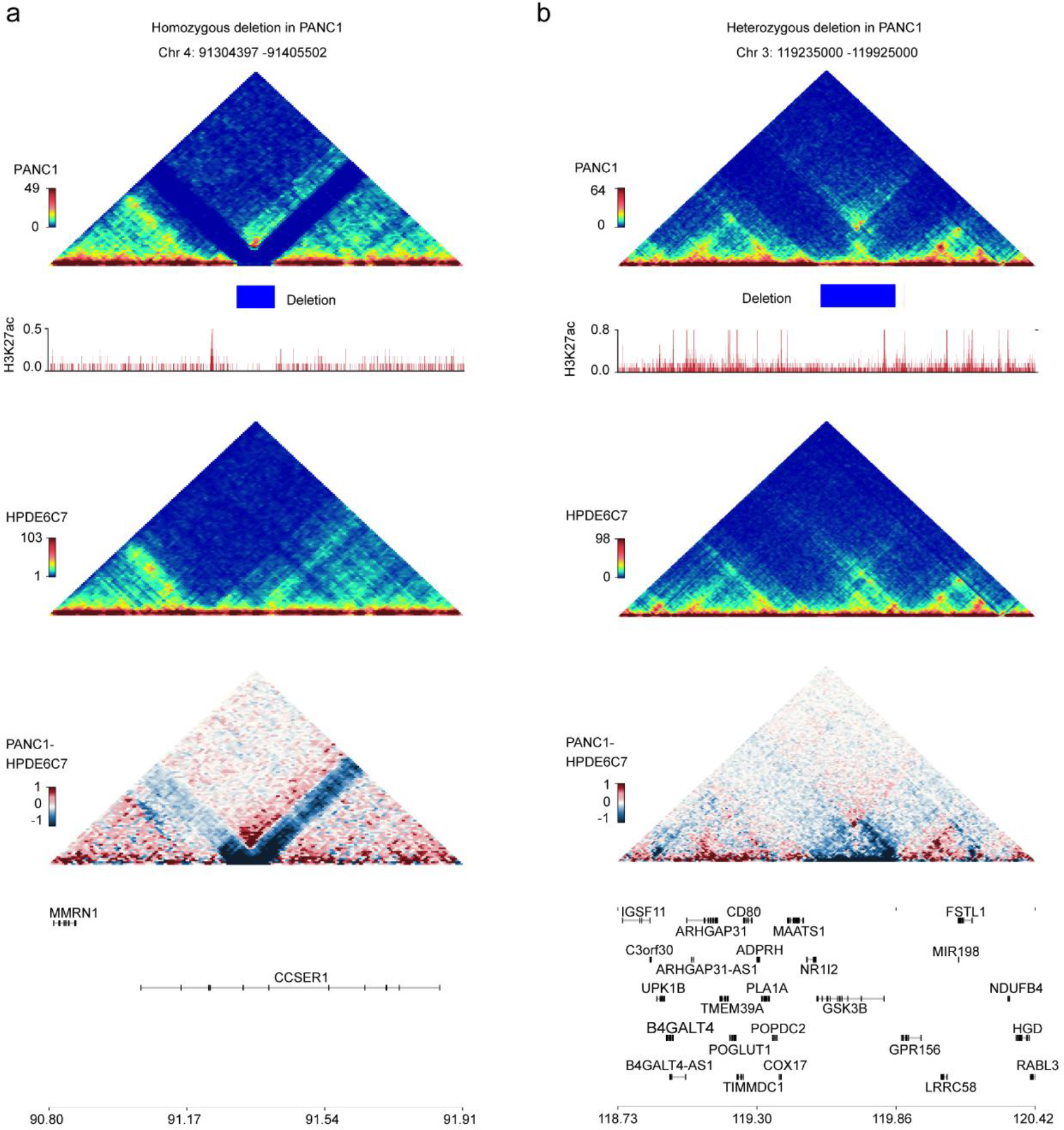
Examples of the impact of cross-CDB deletion purity on the chromatin folding domain in PANC1. Triangle heatmaps represent chromatin contact frequency, with the top showing PANC1, middle showing HPDE6C7, and bottom showing the subtractive results. Histogram representing roadmap epigenome enhancer activity, marked by H3K27ac, in PANC1 (red). a, Homozygous cross-CDB deletion is associated with CD fusion. b, No significant enhancement of adjacent CD interactions is observed in the heterozygous cross-CDB deletion.

**Supplementary Fig. 7.**
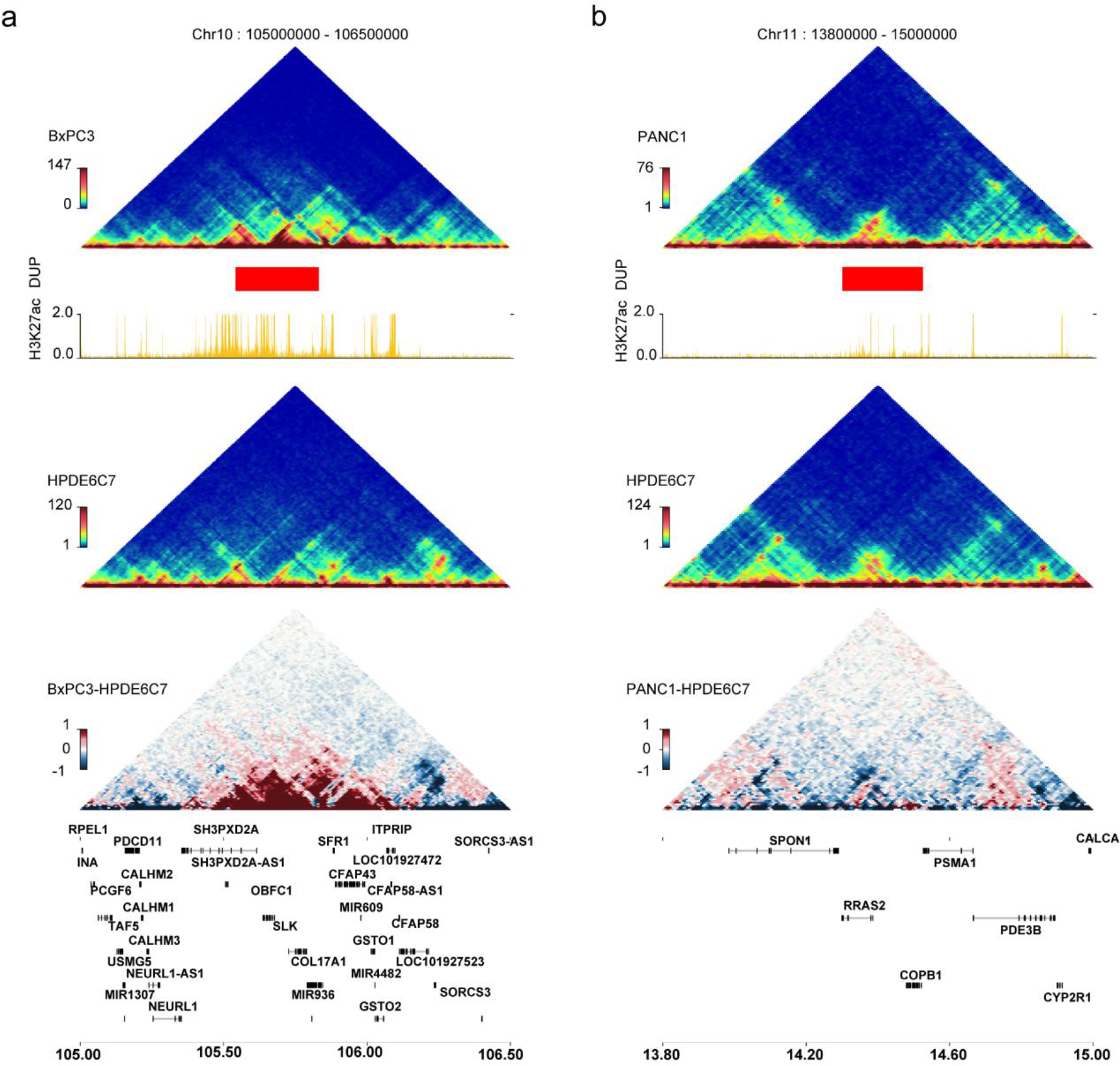
Examples of the impact of cross-CDB duplication purity on the chromatin folding domain in BxPC3 and PANC1. Triangle heatmaps represent chromatin contact frequency, with the top showing cancer cells, middle showing HPDE6C7, and bottom showing the subtractive results. The histogram represents roadmap epigenome enhancer activity, marked by H3K27ac, in cancer cells (red). a, Homozygous cross-CDB duplication is associated with neo-CD formation in the BxPC3 cell line. b, No significant enhancement of adjacent CD interactions was observed in the heterozygous cross-CDB duplication in the PANC1 cell line.

**Supplementary Fig. 8.**
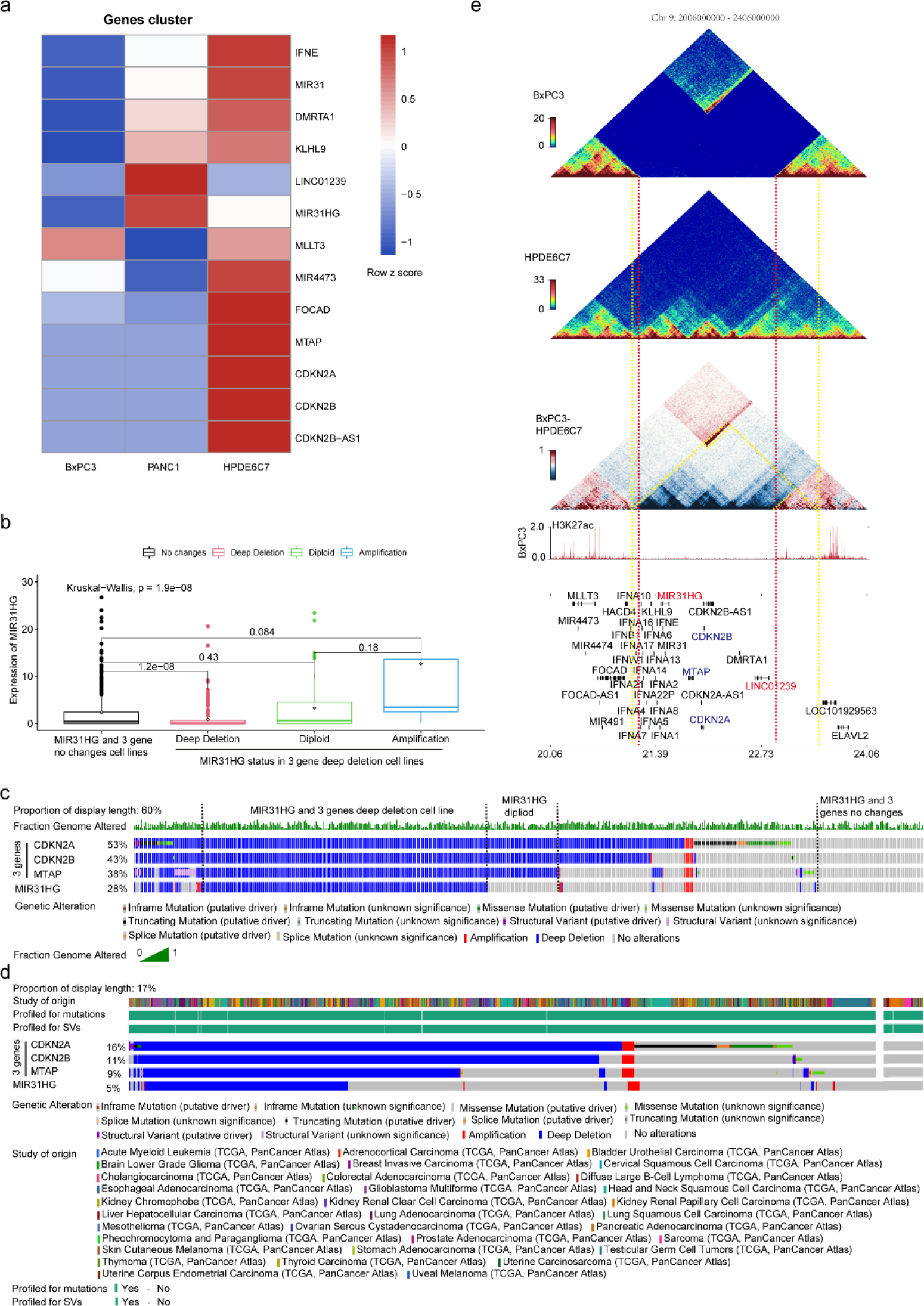
CDKN2A homozygous deletion is associated with MIR31HG upregulation in part through concomitant adjacent genome amplification and CD fusion. **a**, Heatmap representation of RNA-seq results for genes in the CDKN2A-CDKN2B-MTAP deletion region corresponding to Fig. 5a. The heatmap shows the row z score of FPKM normalized read counts. **b**, MIR31HG expression levels in cases of different mutation states of MIR31HG and the three-gene deletion (CDKN2A, CDKN2B and MTAP) in pancancer cell lines from CCLE. P-value is obtained by Kruskal–Wallis test. **c**, Genetic alteration display of CDKN2A, CDKN2B, MTAP and MIR31HG in 807 cancer cell samples from the CCLE dataset by CBioPortal (see URLs). Limited by page size, only 60% of the primary diagram is presented; this includes all cancer cell lines with deep deletion of MIR31HG and the three genes. **d**, Genetic alterations in CDKN2A, CDKN2B, MTAP and MIR31HG in 8359 pancancer samples from TCGA dataset by CBioPortal (see URLs). Limited by page size, only 17% of the primary diagram is presented; this includes all cancer cell lines with deep deletion of MIR31HG and the three genes. **e**, Diagram showing the impacts of CDKN2A homozygous deletion on 3D chromatin folding domains in BxPC3. Triangle heatmaps represent chromatin contact frequency, with the top showing BxPC3, middle showing HPDE6C7, and bottom showing the subtractive results. The histogram below represents roadmap epigenome enhancer activity, marked by H3K27ac, in BxPC3. The red dashed line denotes the break points of the homozygous deletion, and the yellow dashed line marks the boundaries of the fused CD. The yellow dashed line in the bottom triangle heatmap indicates the enhanced interactions of the adjacent CD.

**Supplementary Fig. 9.**
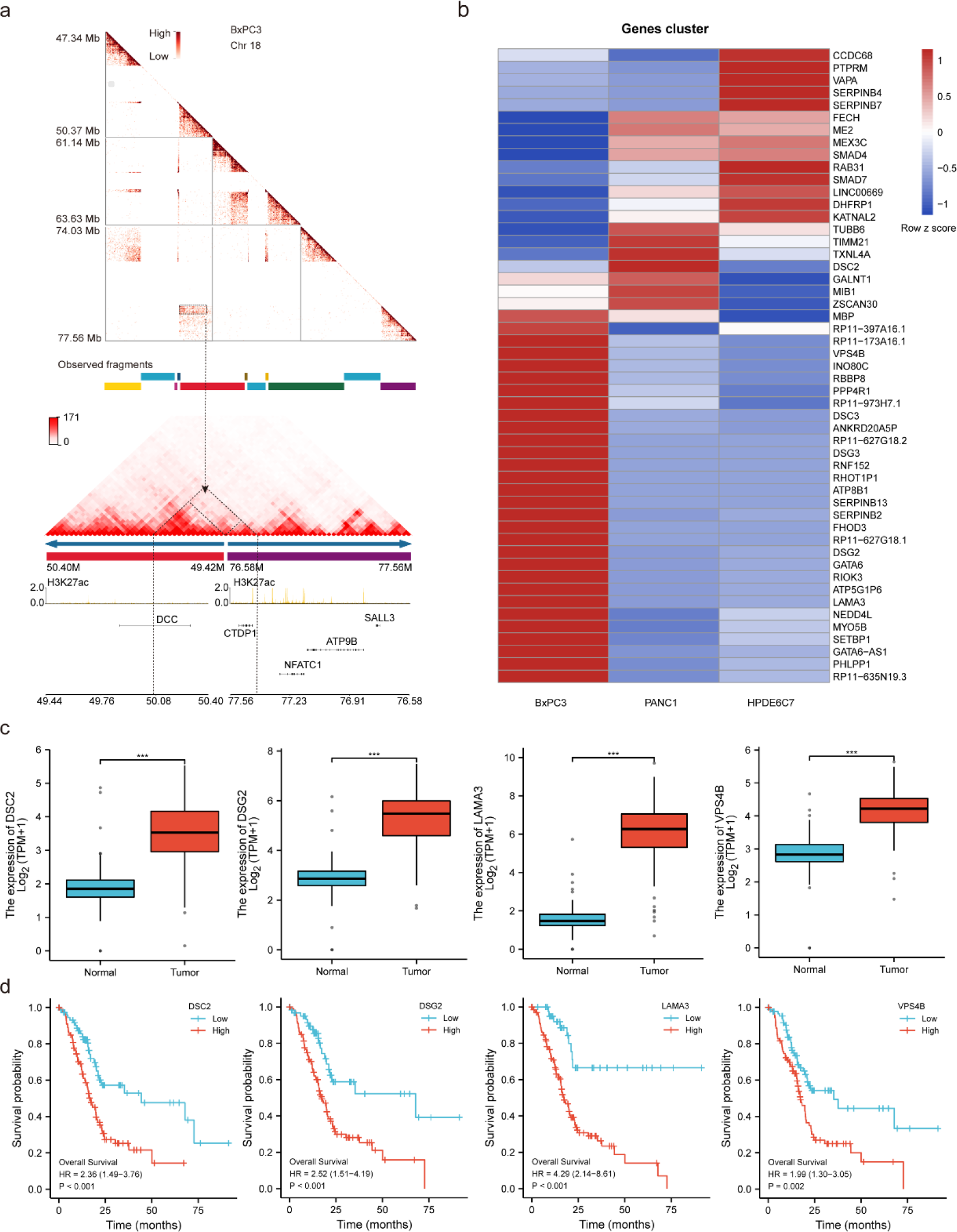
Identification of complex genomic rearrangement associated with SMAD4 deletion in PDAC. **a**, Enlarged heatmap indicated by triangles and rectangles in Fig. 6c. The dashed box shows an example of the aberrant enhanced interaction (neo-CD) in the junction region of rearranged purple and red genome fragments in chromosome 18 of BxPC3 by NeoLoopFinder. The dashed triangle denotes the neo-CD corresponding to ectopic interactions in the enlarged heatmap with the dashed black arrow indicated. The histograms below represent roadmap epigenome enhancer activity, marked by H3K27ac, in BxPC3 (yellow). **b**, Heatmap representation of RNA-seq results for differentially expressed gene-associated complex rearrangements in chromosome 18 in BxPC3. The heatmap shows the row z score of FPKM normalized read counts. **c**, Expression of DSC2, DSG2, LAMA3, and VPS4B in pancreatic cancer and normal control tissues from TCGA and GTEx (n = 350). P-values were obtained by Wilcoxon rank-sum test. ***p ≤ 0.001. **d**, Kaplan–Meier curves for overall survival according to DSC2, DSG2, LAMA3, and VPS4B expression in the TCGA pancreatic cancer dataset (n = 178). P values were obtained by Cox regression via the R package (version 3.6.3).

### Supplementary Tables are attached to supplementary table files.

Supplementary Table 1 | Quality control of long-read sequencing data in HPDE6C7, PANC1 and BxPC3.

Supplementary Table 2 | The number of different types of SVs detected by long-read sequencing in HPDE6C7, PANC1 and BxPC3.

Supplementary Table 3 | The number of different types of cancer-specific SVs detected by long-read sequencing in PANC1 and BxPC3 cells.

Supplementary Table 4 | List of genes directly affected by SVs in exon regions of the genome in HPDE6C7, PANC1 and BxPC3.

Supplementary Table 5 | List of cancer-specific genes directly affected by SVs in exon regions of the genome in PANC1 and BxPC3.

Supplementary Table 6 | List of cancer-specific human cancer genes directly affected by SVs in exon regions of the genome in PANC1 and BxPC3.

Supplementary Table 7 | Comparison of HPDE6C7, PANC1 and BxPC3 by Cellosaurus.

Supplementary Table 8 | Quality control of Hi-C sequencing data.

Supplementary Table 9 | A/B transition in BxPC3 and PANC1 compared with HPDE6C7.

Supplementary Table 10 | Significantly differentially expressed genes in the A/B transition (Log2FC > 1 and q < 0.05).

Supplementary Table 11 | The number and length of CDs called by HiCDB in HPDE6C7, PANC1 and BxPC3 cells.

Supplementary Table 12 | The number and percentage of CDBs called by HiCDB in HPDE6C7, PANC1 and BxPC3 cells.

Supplementary Table 13 | Gene list of specific CDs of PANC1 and BxPC3.

Supplementary Table 14 | Specific loops in PANC1 and BxPC3.

Supplementary Table 15 | The distribution of cancer-specific SVs in the A/B compartments of PANC1.

Supplementary Table 16 | The distribution of cancer-specific SVs in the A/B compartments of BxPC3.

Supplementary Table 17 | The distribution of cancer-specific SVs in CDs of BxPC3 and PANC1.

Supplementary Table 18 | Cross-CDB-Del and CD fusion in BxPC3 and PANC1.

Supplementary Table 19 | Gene list and expression changes in fused CDs of BxPC3.

Supplementary Table 20 | Gene list and expression changes in fused CDs of PANC1.

Supplementary Table 21 | Gene expression change in the Chr 9 CDKN2A-CDKN2B- MTAP deletion region.

Supplementary Table 22 | MIR31HG expression in the CCLE database.

Supplementary Table 23 | MIR31HG expression in TCGA database.

Supplementary Table 24 | The expression level of three genes in the neo-CD of the rearranged fragment junction region.

Supplementary Table 25 | List of differentially expressed genes in chromosome 18.

